# A lipopeptidomimetic of transcriptional activation domains selectively disrupts Med25 PPIs

**DOI:** 10.1101/2023.03.24.534168

**Authors:** Olivia N. Pattelli, Estefanía Martínez Valdivia, Matthew S. Beyersdorf, Clint S. Regan, Mónica Rivas, Sofia D. Merajver, Tomasz Cierpicki, Anna K. Mapp

## Abstract

Short amphipathic peptides are capable of binding to transcriptional coactivators, often targeting the same binding surfaces as native transcriptional activation domains. However, they do so with modest affinity and generally poor selectivity, limiting their utility as synthetic modulators. Here we show that incorporation of a medium-chain, branched fatty acid to the N-terminus of one such heptameric lipopeptidomimetic (34913-8) increases the affinity for the coactivator Med25 >10-fold (*Ki* >>100 μM to 10 μM). Importantly, the selectivity of 34913-8 for Med25 compared to other coactivators is excellent. 34913-8 engages Med25 through interaction with the H2 face of its <u>Ac</u>tivator <u>I</u>nteraction <u>D</u>omain and in doing so stabilizes full-length protein in the cellular proteome. Further, genes regulated by Med25-activator PPIs are inhibited in a cell model of triple-negative breast cancer. Thus, 34913-8 is a useful tool for studying Med25 and the Mediator complex biology and the results indicate that lipopeptidomimetics may be a robust source of inhibitors for activator-coactivator complexes.

---

During transcription initiation, the transcriptional activation domains (TADs) of activators form protein-protein interactions (PPIs) with coactivator proteins, facilitating assembly of the transcriptional machinery.[1–3] The TADs are often composed of amphipathic sequences bearing a preponderance of acidic (D,E) and hydrophobic (L,V,M,F,W) residues.[4, 5] Many lines of evidence indicate that short amphipathic peptides 7-15 amino acids in length and in either D or L configuration are sufficient to replicate the activity of native transcriptional activation domains when tethered to DNA.[6–11] [12–15] The same short sequences rarely function as effective inhibitors of TAD-coactivator interactions, however, as on their own they exhibit modest affinity and poor selectivity for coactivators.[9,16,17] One successful approach for improving the affinity of short TADs has been to incorporate moieties that induce and/or stabilize helical secondary structures that the peptides are thought to assume upon binding to coactivators.[18,19] This has led, for example, to peptidomimetic inhibitors of CBP/p300 TAZ2-activator complexes that are effective in vitro and in animal models.[20] Nonetheless, given the limited structural data for many activator-coactivator complexes, a structure-independent approach to convert short amphipathic sequences into effective inhibitors would be an important addition to the field.

N-Terminal fatty acid acylation of short peptides has been shown to increase the efficacy, stability, and cell permeability in a variety of applications.[21–26] Kim and co-workers, for example, reported that the palmitoylation of a heptamer discovered in a screen for proliferation PPI modulators produced inhibitors with excellent activity in vitro and in cells.[24] We hypothesized that such a strategy could produce more potent inhibitors of activator-coactivator PPIs, although selectivity for a given coactivator could still be an issue. To test this hypothesis, we chose the coactivator Med25 and its activator PPI network as the target (Figure 1). Med25 is a substoichiometric component of the megadalton Mediator complex and serves to bridge Mediator and transcriptional activators during transcription initiation, including the ETV/PEA3 family (cell migration, motility), ATF6α (stress response), and VP16 (viral infection).[27–33] Several lines of evidence suggest that dysregulation of Med25-activator PPIs, particularly the Med25-ETV/PEA3 network, plays a role in a number of cancers, making Med25 a potentially attractive therapeutic target. [27, 34] Med25 has also emerged as a potential lipid-binding protein, although the underlying biological role is not yet known. [35] It is, however, a highly challenging target with only a single inhibitor reported thus far. [36]

**Figure 1.**
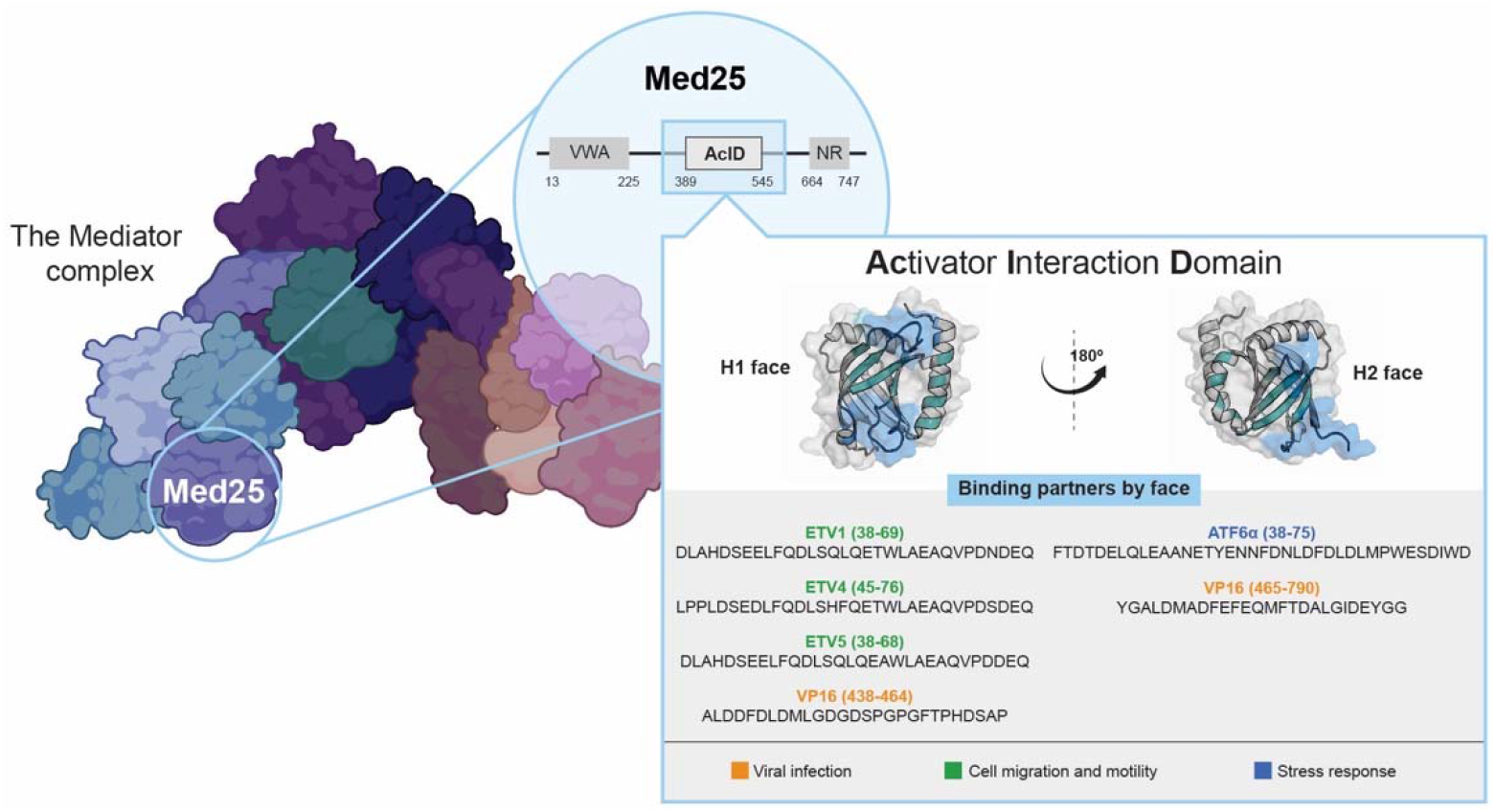
Med25 is a component of the Mediator complex. It uses the two binding surfaces (H1 and H2) of its Activator Interaction Domain (AcID) to complex with amphipathic activators, including the ETV/PEA3 family (ETV1, ETV4, ETV5) VP16, and ATF6α. PDB entry 2XNF was used to generate figure.

Here we demonstrate that incorporation of a branched fatty acid at the N-terminus of a 7-residue peptide leads to >10-fold increase in potency against Med25-activator PPIs in vitro and with excellent selectivity. Further, the lead inhibitor, 34913-8, engages full-length Med25 in the cellular proteome and down-regulates Med25-dependent genes. Taken together, these data suggest that lipopeptidomimetics can be a valuable and thus-far unexplored tool for the molecular intervention of coactivators and their PPI networks.

We chose as a starting point the peptide sequence EDLLLLV, a sequence that contains a balance of hydrophobic and polar/acidic amino acids frequently observed in transcriptional activation domains (Figure 2a). This peptide has a modest affinity for Med25, with a *Ki* of 110 μM against the Med25-ATF6α complex. As reported in Figure 2b, we then tested both the D- and L-variants of the sequences with fatty acid tails of increasing complexity and varied C-termini. The data revealed that the D- and L-versions of the peptide were essentially identical in activity (34913-2 versus 34191-3; 34913-4 versus 34913-5) and thus D-amino acids were used moving forward due to advantageous proteolytic stability. The interchangeability of the D- and L-peptides is consistent with other studies of short amphipathic peptides interacting with coactivators. [9,11, 13] A significant increase in efficacy was observed when a medium chain fatty acid (C11) was added, with 34913-6 and 34913-7 exhibiting low micromolar *K^is^*. However, precipitation was observed during the binding reactions and increasing detergent concentration led to attenuation of inhibitory activity, indicating that 34913-6 and 34913-7 may function as aggregates.[37]

**Figure 2.**
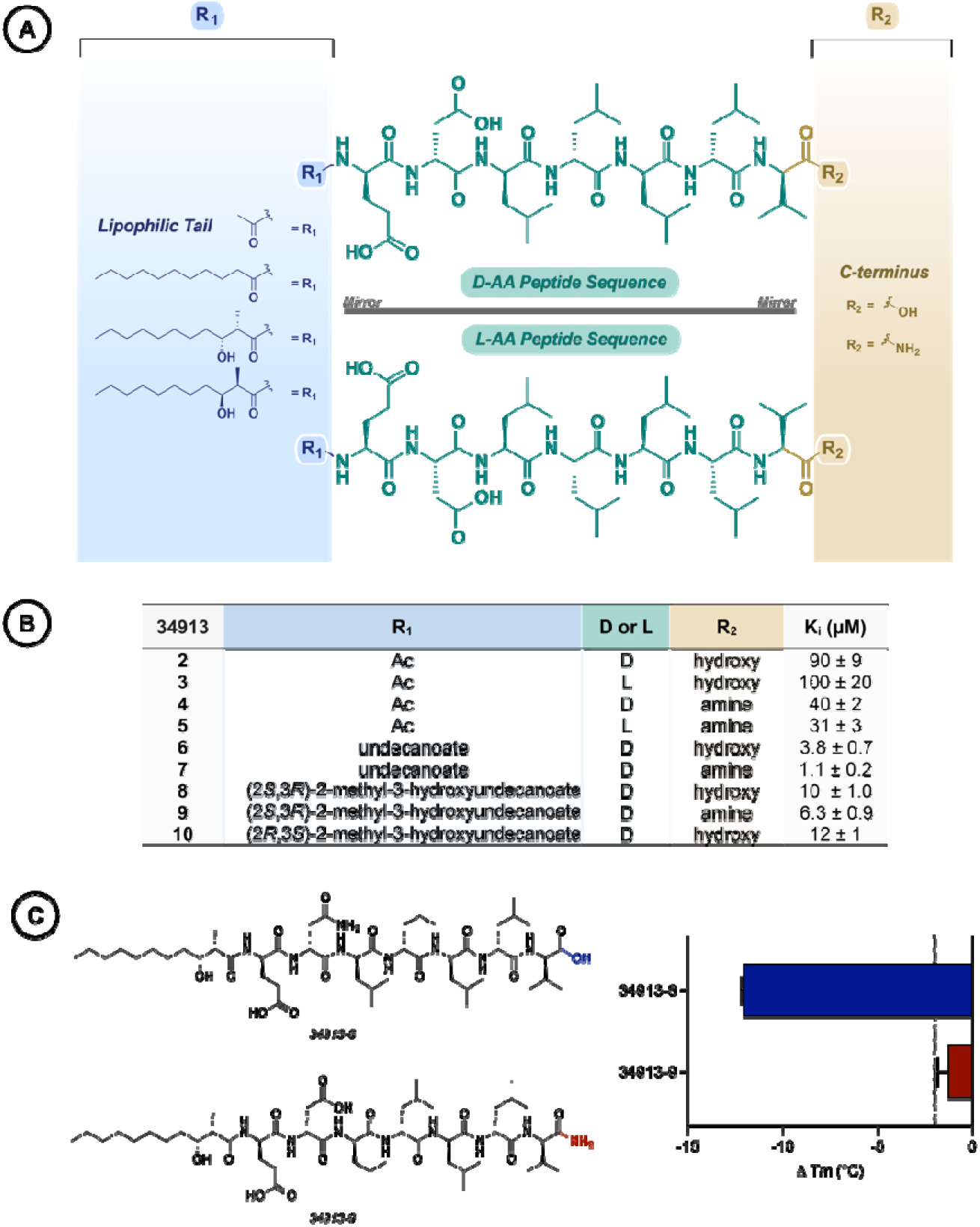
**A)** Schematic of lipopeptidomimetic constructs used in this study consisting of various N-terminal fatty acid acylations, amino acid stereochemistry, and C-termini. B) Inhibition of Med25 AcID•ATF6α (residues 38-75) by lipopeptidomimetics as determined by competitive fluorescence polarization assays. IC_50_ values were measured by titrating the lipopeptidomimetics with 20 nM FITC-ATF6α in complex with Med25 AcID (50% bound). The IC_50_ values were converted to K_i_ values using the apparent K_d_ from direct binding measurements of Med25 AcID•ATF6α and using a published K_i_ calculator. [41] Data shown is the average of three independent experiments performed in technical triplicate with the indicated error (SD). **C)** Change in the melting temperature (ΔT^m^) of 8 μM Med25 AcID in the presence of 5 equiv. of 34913-8 (blue bar) or 34913-9 (red bar), as determined by differential scanning fluorimetry. Temperature-dependent unfolding was monitored using Sypro Orange fluorescence. Values represent the change in melting temperature relative to unbound Med25 AcID control. The ΔT^m^ values are the average of two independent experiments performed in technical triplicate with the indicated error (SD).

A common feature of bacterial lipopeptides is a β-hydroxyl group and, less frequently, α-branching. [38–40] Reasoning that incorporation of such functional groups would increase solubility and reduce aggregation, we incorporated (2S, 3R)-2-methyl-3-hydroxyundecanoate and (2R,3S)-2-methyl-3-hydroxyundecanoate into analogs 34913-8, −9, and −10. The analogs exhibited nearly identical activity, with the stereochemistry having no measurable impact (Figure 2B, 2C). The engagement of 34913-8 and 34913-9 with Med25 was then assessed using differential scanning fluorimetry (DSF) and, notably, incubation of Med25 AcID with 34913-8 induced a significant change in Tm (Figure 2C). In contrast, incubation of Med25 AcID with 34913-9 produced only a minor change in the melting temperature, suggesting a lack of specific engagement of the lipopeptidomimetic with Med25.

As noted earlier, Med25 AcID contains two binding surfaces for amphipathic activators that are allosterically connected. To assess if 34913-8 functions through interaction with one or both of the native binding surfaces, both ^1^H,^15^N- and ^1^H,^13^C-HSQC experiments were carried out (Figure 3). Addition of 1.1 eq of 34913-8 resulted in several ^1^H,^15^N and ^1^H,^13^C-HSQC chemical shift perturbations (CSPs) of residues located predominantly on the H2 binding surface of Med25 AcID. The majority of residues perturbed in the ^1^H,^13^C-HSQC spectra are solvent-exposed and located on the H2 face □-barrel and flanking □-helices (□1 and □2), with the exception of an inward facing side-chain methyl group of L406 located on the □-barrel of the H1 face (Figure 3A).[42] Super-stoichiometric concentrations of 34913-8 showed increased CSPs at both binding faces, though the H1 face lacked notable perturbations of solvent-exposed residues. Taken together, these data suggest that 34913-8 directly binds to the H2 face of Med25 AcID through engagement with the □-barrel and dynamic framing helices and in doing so induces conformational changes.

**Figure 3.**
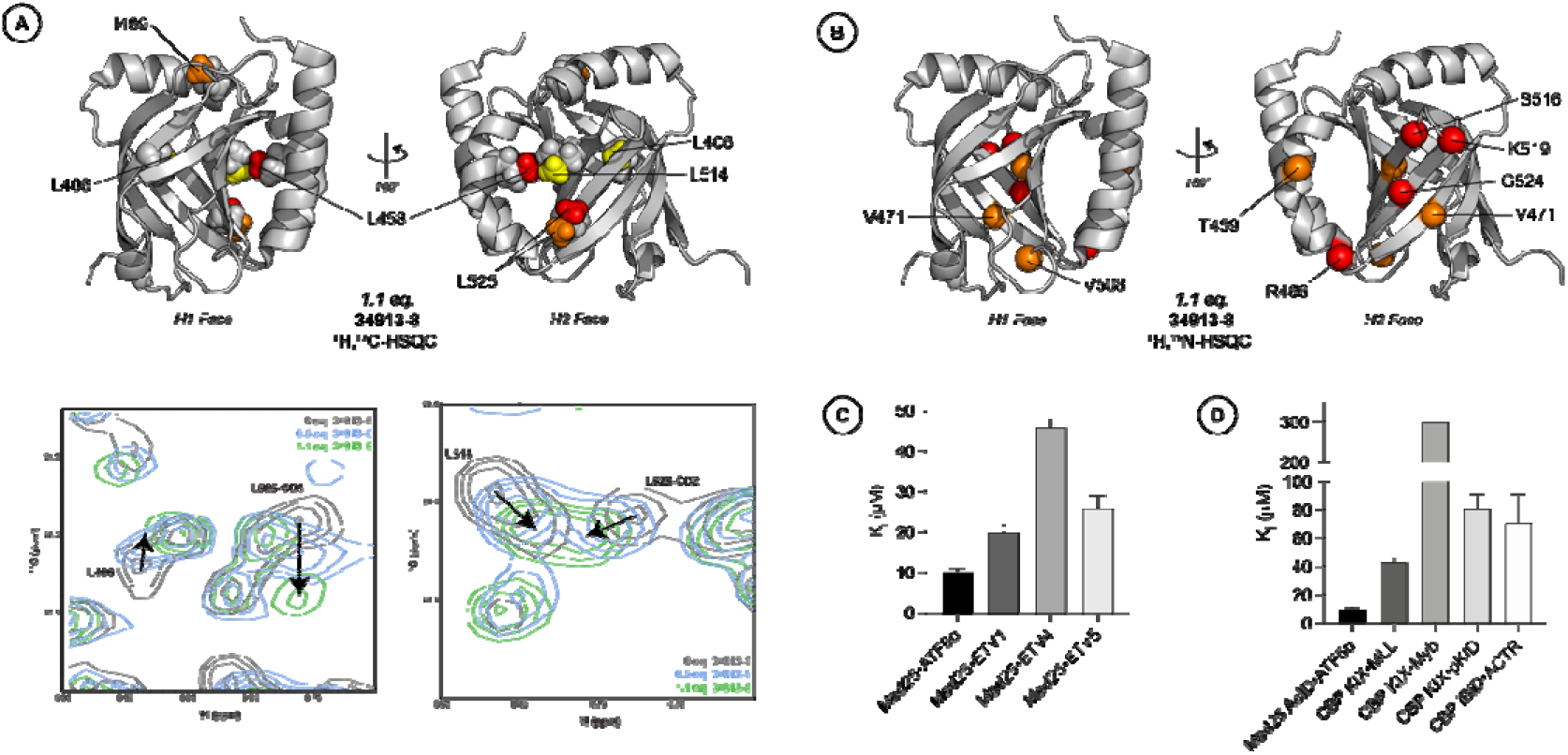
Lipopeptidomimetic 34913-8 is a mixed allosteric/orthosteric inhibitor that engages the H2 face of Med25 AcID. A) ^1^H,^13^C-HSQC CSPS induced by binding of 1.1 equiv. of 34913-8 mapped onto Med25 AcID (PDB ID 2XNF). Yellow = 0.02 ppm - 0.0249 ppm, orange = 0.025 ppm - 0.049 ppm, red ≥ 0.0491. Overlay of ^1^H, ^13^C-HSQC CSPs of free Med25 (dark grey), 0.5 equiv. 34913-8 (light blue) and 1.1 equiv. of 34913.8 (green) for Med25 residues L406, L514, and L525. B) ^1^H, ^15^N-HSQC CSPs induced by binding of 1.1 equiv. of 34913-8 mapped onto Med25 AcID (PDB ID 2XNF). Only residues with CSPs > 0.0851 ppm are labeled. Orange = 0.0851 ppm-0.14 ppm, red ≥ 0. 141. All perturbed residues above signal to noise ratio (≥ 0.02 ppm) found in Supplemental. C) Inhibition of Med25 AcID•ETV/PEA3 PPIs by 34913-8 as determined by competitive fluorescence polarization assays. IC_50_ values were measured by titrating the lipopeptidomimetics against 20 nM FITC-ETV1/ETV4/ETV5 in complex with Med25 AcID (50% bound). The IC_50_ values were converted to K_i_ values using the apparent K_d_ from direct binding measurements of Med25 AcID•ETV/PEA3 PPIs using a published K_i_ calculator. [41] Data shown is the average of three independent experiments performed in technical triplicate with the indicated error (SD). D) Selectivity of 34813-8 for Med25 AcID as determined by the inhibition of related PPI networks using competitive fluorescence polarization assays. IC_50_ values using a suite of coactivators bound to FITC-activators at 20 nM (CBP KIX•MLL/Myb/pKID, CBP IBiD• ACTR). The IC_50_ values were converted to K_i_ values using a published K_i_ calculator [41] and the corresponding K_d_ value of each coactivator•activator direct binding measurement. Med25 AcID•ATF6α and CBP KIX•MLL/Myb data shown is the average of three independent experiments performed in technical triplicate with the indicated error (SD). CBP KIX•pKID and CBP IBiD• ACTR data shown is the average of two independent experiments performed in technical triplicate with the indicated error (SD). No error bars are shown for the IC_50_ against CBP KIX•Myb because the IC_50_ was greater than the highest concentration of 34913-8 tested (300 μM), and thus, we can accurately report the IC_50_ only as > 300 μM.

In earlier work we showed that endogenous activators and small-molecule modulators of Med25 engage dynamic substructures adjacent to the β-barrel surfaces and in doing so effect conformational changes and allosteric communication between the H1 and H2 faces.[42, 43] Even in the case of activators ETV1, ETV4, and ETV5 that interact with the same H1 surface, small sequence differences lead to engagement with distinct dynamic substructures and correspondingly distinct conformational changes of the activator-Med25 complexes. [42] Because 34913-8 engages with the flexible framing helices of the H2 face and appears to stimulate broader conformational changes, we hypothesized that allosteric effects on H1 binding activators should occur. Consistent with the HSQC NMR results, 34913-8 inhibits the Med25 H1-binding ETV/PEA3 family of transcription activators, ETV1, ETV4, and ETV5 (Figure 3C). Further, consistent with each of the ETV/PEA3 activators interacting with the H1 surface in distinct modes, 34913-8 demonstrates ~2-fold selectivity for Med25•ETV1/5 compared to Med25•ETV4. We have previously shown that engagement of dynamic substructures within coactivators such as Med25 not only produces allosteric inhibitors but also inhibitors with increased selectivity for their cognate coactivator.[36] The selectivity of 34913-8 for Med25 relative to a broader range of coactivators was assessed in a series of competitive inhibition essays (Figure 3D). 34913-8 showed excellent selectivity, with >6-fold preference for Med25 across the range of coactivators tested.

Lipopeptidomimetic *34913-8* engages not just with the isolated AcID motif but also with fulllength Med25. To test this, we utilized the triple negative breast cancer cell line VARI068 that, as previously described, exhibits upregulated Med25 and its cognate ETV/PEA3 activator binding partners. [36] Incubation of freshly prepared VARI068 nuclear extracts with 34913-8 lead to stabilization of endogenous Med25, indicating target engagement of AcID in the context of fulllength Med25 (Figure 4a). Consistent with the previous observations, incubation with 34913-9 did not induce stabilization of endogenous Med25. The Med25•ETV/PEA3 PPIs are implicated in cell proliferation, migration, and invasion pathways that are frequently associated with carcinogenesis. For this reason, the gene transcripts of the Med25•ETV5-regulated protein, MMP2 were quantified with qPCR using 34913-8. Treatment with 34913-8 downregulated MMP2 gene expression, while treatment with negative control 34913-9 did not (Figure 4B, right). Collectively, this data is consistent with 34913-8 engaging Med25 in cells and altering its PPI network, resulting in downstream effects on gene expression, while analog 34913-9 is unable to specifically engage Med25.

**Figure 4.**
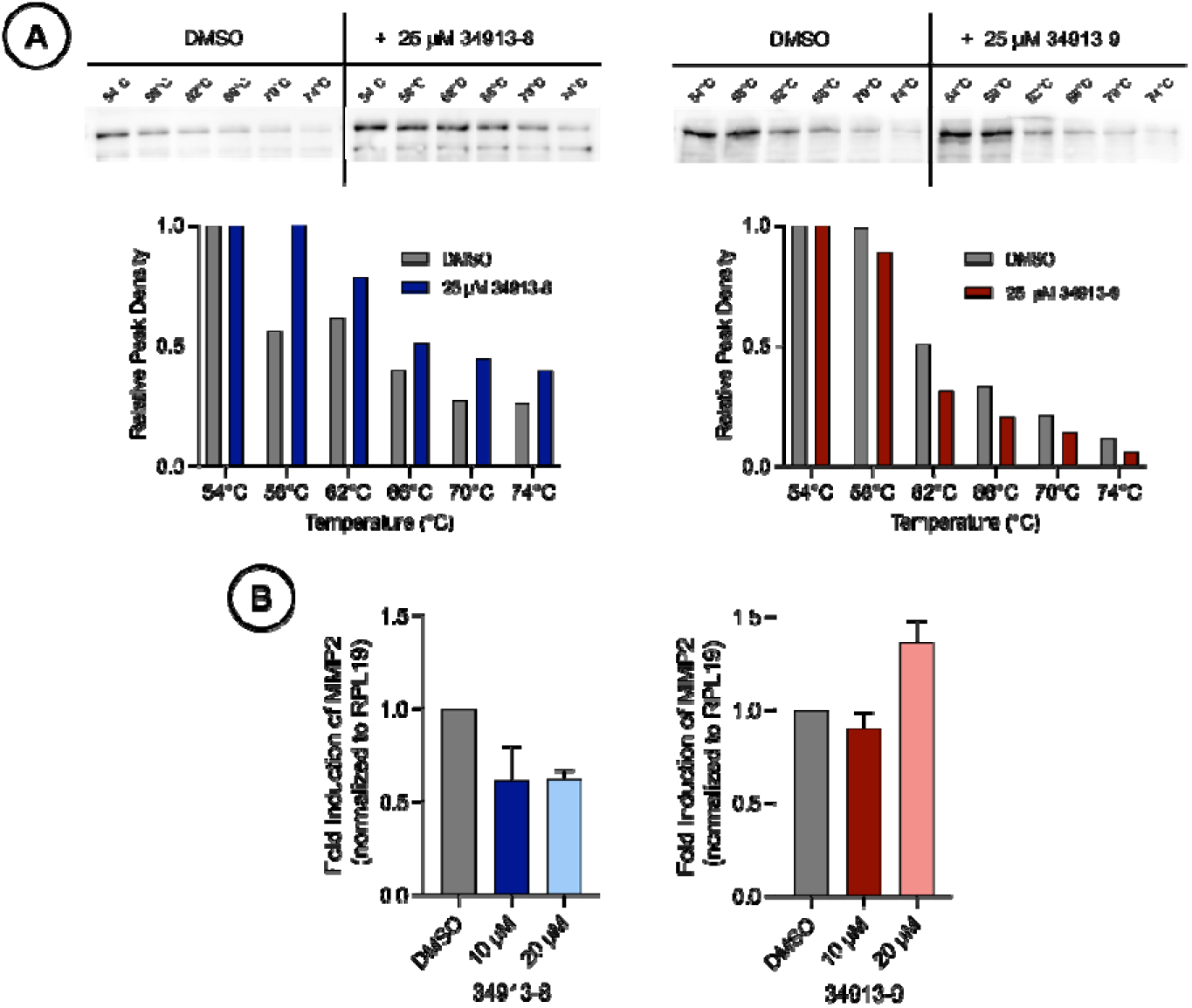
A) Lipopeptidomimetic 34913-8 stabilizes full length Med25 in VARI068 cell extracts. Cellular thermal shift assays (CETSA) were performed by dosing VARI068 nuclear extracts with 25 μM 34913-8, 34913-9, or equivalent DMSO and subjecting the samples to a range of temperahires. Western blots using a Med25 antibody show an increased band intensity in 34913-8-dosed samples compared to the control 34913-9, indicating thermal shift stabilization and target engagement. Data in the bar graph is normalized to the DMSO control (grey) that is equal to 1. Data shown is representative of experiments performed in biological duplicates. B) Lipopeptidomimetic 34913-8 shows inhibition of Med25 in a cellular context. Analysis and quantification of the transcript levels of the Med25•ETV5-dependent gene MMP2 in the triple-negative breast cancer cell line VARI068 was performed using qPCR. Results indicate that treatment with increasing concentrations of 34913-8 results in the decrease of MMP2 transcript levels. By comparison, increasing concentrations of 34913-9 do not yield changes in MMP2 transcript levels. Values are normalized to the reference gene RPL19. Data shown is the average of three independent experiments performed in technical triplicate with the indicated error (SD).

Lipopeptidomimetic 34913-8 represents the first reversible inhibitor that targets the Med25•activator PPIs, with the lipid tail contributing significantly to its activity. The peptide sequence of 34913-8 displays considerable similarity to amphipathic activators, sequences that on their own are typically quite promiscuous binders. In contrast, 34913-8 shows excellent selectivity for Med25 AcID (>6-fold). Taken together the data indicate that the linear lipopeptide architecture of 34913-8 is an excellent starting point for the development of a broader array of coactivator modulators. Future work includes assessing linear lipopeptide natural products as well as synthetic lipopeptidomimetics differing in both amino acid sequence and lipid architectures.

## Acknowledgments

The authors greatly appreciate colleagues Prof. D.H. Sherman, Dr. A. Tripathi, and P. Schulz, who inspired this work by sharing the unpublished structure of a novel bacterial lipopeptide similar in architecture. Thank you to F. Gu for synthesis assistance. The authors acknowledge financial support from the National Institutes of Health (CA242018 and GM136356 to A.K.M.; NIH-P30CA046592 to the Rogel Cancer Center Core) and and the Breast Cancer Research Foundation (to SDM).

