## Supporting information for "A lipopeptidomimetic of transcriptional activation domains selectively disrupts Med25 PPIs"

### Experimental Procedures

*Protein Expression and Purification.* The Med25 expression plasmid pET21b-Med25(394-543)-His6 was generously provided by Professor Patrick Cramer of the Max Planck Institute for Biophysical Chemistry, Göttingen, Germany.[1] WT Med25 AcID was expressed as previously described with slight modifications.[1-2] Plasmid was transformed into heat-shock competent E. coli BL21 (DE3) cells were grown in TB Broth supplemented with 0.1 mg/mL of ampicillin. $^{13}C,^{15}N$-labeled Med25 AcID was expressed as previously described with slight modifications. [1-2] Plasmid was transformed into heat-shock competent E. coli BL21 (DE3) cells and cells were grown in M9 minimal media supplemented with 0.1 mg/mL of ampicillin, 1 g/L $^{15}NH_4Cl$, 2 g/L $^{13}C$-D-glucose, and 0.5% $^{13}C,^{15}N$-labeled Bioexpress media (all labeled components were purchased from Cambridge Isotopes). Purification of both proteins was performed according to previously described methods using an AKTA Pure FPLC.[1-2] Fractions containing protein were dialyzed into storage buffer (10 mM sodium phosphate, 100 mM NaCl, 10% glycerol, pH 6.8) or NMR buffer (20 mM sodium phosphate, 150 mM NaCl, pH 6.5). Concentration was determined via ultraviolet/visible (UV/Vis) spectroscopy on a NanoDrop instrument at 280 nm using an extinction coefficient of 22,460 $M^{-1}cm^{-1}$. Protein was aliquoted, flash frozen in liquid $N_2$ and stored at –80 ºC until further use. Protein identity was confirmed by mass spectrometry (Agilent 6545 LC/Q-TOF) and purity was assessed via sodium dodecyl sulphate–polyacrylamide gel electrophoresis (SDS-PAGE) on a 4-12% bis-tris gel stained using Quick Coomassie (Anatrace).

CBP KIX (586-672) was expressed as previously described with slight modifications.[3] Plasmid was transformed into heat-shock competent E. coli BL21 (DE3) cells and grown in TB Broth supplemented with

0.1 mg/mL of ampicillin. Protein purification was performed according to previously described methods using an AKTA Pure FPLC. [3] Cell pellets were resuspended in ~35 mL lysis buffer (10 mM phosphate, 300 mM NaCl, 10 mM imidazole, pH 7.2), 40 μL β-mercaptoethanol, and a cOmplete protease inhibitor tablet (Roche). Cells were lysed by sonication and insoluble cellular material was pelleted by centrifugation at 9500 rpm for 20 min at 4 ºC. The lysate filtered and loaded onto an AKTA Pure FPLC equipped with a 5 mL Ni HisTrap HP column (GE Healthcare) pre-equilibrated with wash buffer (10 mM phosphate, 300 mM NaCl, 10 mM imidazole, pH 7.2). CBP KIX was then purified using a gradient of 10–600 mM imidazole (other buffer components were constant), and fractions containing CBP KIX were pooled and subjected to secondary purification using a HiTrap SP HP cation exchange column (GE Healthcare) using buffer (50 mM sodium phosphate, 1 mM dithiothreitol (DTT), pH 7.2) with gradient of 0-1 M NaCl. Fractions containing pure protein were dialyzed into storage buffer (10 mM sodium phosphate, 100 mM NaCl, 10% glycerol, pH 6.8) overnight at 4 ºC. Concentration was determined via ultraviolet/visible (UV/Vis) spectroscopy on a NanoDrop instrument at 280 nm using an extinction coefficient of 12,950 $M^{-1}cm^{-1}$. Aliquots were flash frozen in liquid $N_2$ and stored at –80 ºC until further use. Protein identity was confirmed by mass spectrometry (Agilent 6545 LC/Q-TOF) and purity was assessed via sodium dodecyl sulphate–polyacrylamide gel electrophoresis (SDS-PAGE) on a 4-12% bis-tris gel stained using Quick Coomassie (Anatrace).

*Lipophilic Tail Synthesis*

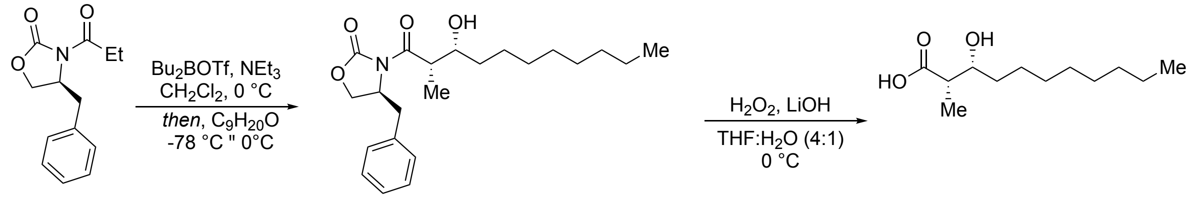


(*S*)-**S1** **I** **S2** **31% Yield**

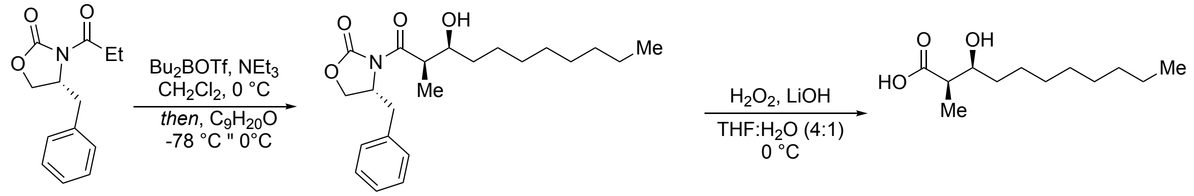


(*R*)-**S3** **II** **S4** **31% Yield**

(2*S*, 3*R*)-2-methyl-3-hydroxyundecanoic acid (**S2**) and (2*R*, 3*S*)-2-methyl-3-hydroxyundecanoic acid (**S4**) were synthesized according to standard aldol reaction protocols.[4] (*2R, 3R)*-2-Methyl-3-hydroxyundecanoic acid has been previously described as has racemic 2-methyl-3-hydroxyundecanoic acid.[5] The spectral data for **S2** and **S4** are provided below and in SI figures S1-S4.

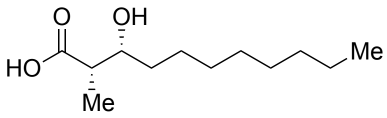


(2*S*,3*R*)-3-hydroxy-2-methylundecanoic acid (**S2**) $R_f$ (4:1 hexanes: ethyl acetate v/v): 0.12. $^{1}$H NMR (599 MHz, $CDCl_3$) δ 1H NMR (599 MHz, $CDCl_3$) δ 3.96 – 3.90 (m, 1H), 2.62 – 2.53 (m, 1H), 1.57 – 1.35 (m, 3H), 1.33 – 1.21 (m, 11H), 1.19 (d, *J* = 7.3 Hz, 3H), 0.86 (t, *J* = 7.0 Hz, 3H). $^{13}$C NMR (151 MHz, $CDCl_3$) δ 181.31, 71.73, 44.04, 33.64, 31.84, 29.51, 29.49, 29.23, 25.97, 22.64, 14.08, 10.36.

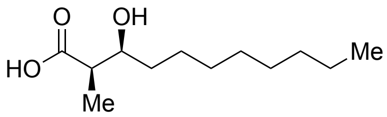

(2*R*,3*S*)-3-hydroxy-2-methylundecanoic acid (**S4**) $R_f$ (4:1 hexanes: ethyl acetate v/v): 0.12. $^1$H NMR (599 MHz, $CDCl_3$) δ 3.93 (dt, *J* = 8.5, 3.8 Hz, 1H), 2.63 – 2.55 (m, 1H), 1.51 – 1.38 (m, 3H), 1.34 – 1.23 (m, 11H), 1.19 (d, *J* = 7.2 Hz, 3H), 0.86 (t, *J* = 6.7 Hz, 3H). $^{13}$C NMR (151 MHz, $CDCl_3$) δ 180.82, 71.73, 44.04, 33.63, 31.84, 29.51, 29.49, 29.23, 25.97, 22.65, 14.09, 10.38.

*Solid-Phase Synthesis and HPLC Purification of Activator Peptides*

Activator peptides were synthesized using standard fluorenylmethyloxycarbonyl (FMOC) solid-phase synthesis methods on a Liberty Blue Microwave Synthesizer (CEM). Fmoc-protected β-alanine or Fmoc-protected 8-amino-3,6-dioxaoctanoic acid (AEEAc) linker was added to the N-terminus of all labelled peptides prior to fluorescent labeling. Resin was deprotected using 20% piperidine in DMF. Peptides were then treated with 1.5 equiv. fluorescein isothiocyanate (FITC, ThermoFisher) in the presence of 5% N,N-diisopropylethylamine (DIPEA, Sigma Aldrich) in DMF for 15 hr. IBiD activator peptide was acetylated at the N-terminus using a cocktail of acetic anhydride and triethylamine (TEA, Fisher Scientific) in DCM for 30 min. All activator peptides were cleaved from resin using a 95% trifluoroacetic acid (TFA, Sigma Aldrich), 2.5% triisopropylsilane (TIPS, Sigma Aldrich), and 2.5% $H_2O$ solution for two hours. Resin was filtered, then solution was concentrated and precipitated in cold diethyl ether.

Precipitated peptide was dissolved in 1:1 solution of 0.1% TFA in water and acetonitrile with minimal ammonium hydroxide, syringe-filtered, and purified via high-performance liquid chromatography (HPLC) purification. Peptides were purified via an Agilent 1260 Series HPLC using a C18 column (Phenomenex) and a 10-50 gradient elution of acetonitrile and 0.1% TFA in water over 40 min. Pure fractions were pooled, lyophilized, and stored at -20 ºC until further use. Identity and purity of peptides were confirmed using analytical HPLC and mass spectrometry under negative ion mode (Agilent 6545 LC/Q-TOF). FITC-labelled pure peptides were dissolved in minimal dimethyl sulfoxide (DMSO) and quantified on a NanoDrop instrument at 495 nm using extinction coefficient 72,000 $M^{-1}cm^{-1}$. Acetylated IBiD was dissolved in minimal dimethyl sulfoxide (DMSO) and quantified on a NanoDrop instrument at 280 nm using extinction coefficient 1,490 $M^{-1}cm^{-1}$.

| Peptide | Sequence (N'-C') |
| --- | --- |
| *ATF6α (38-75)* | FITC-βAla-FTDTDELQLEAANETYENNFDNLDFDLDLMPWESDIWD |
| *ETV1 (38-69)* | FITC-βAla-DLAHDSEELFQDLSQLQETWLAEAQVPDNDEQ |
| *ETV4 (45-76)* | FITC-βAla-LPPLDSEDLFQDLSHFQETWLAEAQVPDSDEQ |
| *ETV5 (38-68)* | FITC-βAla-DLAHDSEELFQDLSQLQEAWLAEAQVPDDEQ |
| *MLL (2840-2858)* | FITC-βAla-DCGNILPSDIMDFVLKNTP |
| *Myb (219-316)* | FITC-βAla-KEKRIKELELLLMSTENELKGQQVLP |
| *ACTR (1041-1088)* | FITC-βAla-PSNLEGQSDERALLDQLHTLLSNTDATGLEEIDRALGIPELVNQGQAL |
| *pKID (105-133)* | FITC-(AEEAc)-TDSQKRREILSRRPS($PO_4$)YRKILNDLSSDAPG |
| *IBiD (2063-2111)* | Ac-SPSALQDLLRTLKSPSSPQQQQQVLNILKSNPQLMAAFIKQRTAKYVAN |

*Table S1***.** Sequences of activator peptides used in this study.

*Solid-Phase Synthesis and HPLC Purification of Lipopeptidomimetics*

Peptides of lipopeptidomimetics were synthesized using Rink Amide Protide resin (CEM) or Cl-MPA Protide resin (CEM). Rink Amide Protide resin was deprotected for 30 min. using 20% piperidine in DMF and the first amino acid (5 equiv, ChemImpex and NovaBiochem) was coupled using 1:1:1 ratio of 0.5 M hexafluorophosphate benzotriazole tetramethyl uronium (HBTU) in DMF, 0.49 M hydroxybenzotriazole (HOBT) in DMF, and 1.0 M DIPEA in DMF for 1.5 hrs. Cl-MPA Protide resin was coupled with the first amino acid (5 equiv, ChemImpex and NovaBiochem) three times using 1:1 ratio of 0.125 M KI in DMF and 1.0 M DIPEA in DMF for 1 hr. each. After first residue was coupled, resins were rinsed 3x with DMF, deprotected for 30 min. with 20% piperidine in DMF, and then coupled with next amino acid for 1.5 hrs. This was repeated until the peptide sequence was complete. A final deprotection was performed and analogs 34913-2 through 34913-5 were acetylated at the N-terminus using a cocktail of acetic anhydride and triethylamine (TEA, Fisher Scientific) in DCM for 30 min. Analogs 34913-6 and 34913-7 were coupled with undecanoic acid (5 equiv.) using 1:1:1 ratio of 0.5 M HBTU in DMF, 0.49 M HOBT in DMF, and 1.0 M DPIEA in DMF for 15 hr. Analogs 34913-8 and 34913-9 were coupled to crude (2S, 3R)-2-methyl-3hydroxyundecanoic acid (5 equiv.) using 1:1:1 ratio 0.5 M HBTU in DMF, 0.49 M HOBT in DMF, and 1.0 M DIPEA in DMF for 15 hr. Analog 34913-10 was coupled to crude (2R, 3S)-2-methyl-3-hydroxyundecanoic acid (2 equiv.) using 1:1:1 ratio 0.5 M HBTU in DMF, 0.49 M HOBT in DMF, and 1.0 M DIPEA in DMF for 15 hr.

All lipopeptides were cleaved from resin using a 95% trifluoroacetic acid (TFA, Sigma Aldrich), 2.5% triisopropylsilane (TIPS, Sigma Aldrich), and 2.5% $H_2O$ solution for two hrs. Resin was filtered, then solution was concentrated and precipitated in cold diethyl ether. The precipitated lipopeptide was dissolved in MeOH, filtered, and purified via high-performance liquid chromatography (HPLC) purification. Peptides were purified via an Agilent 1260 Series HPLC using a C18 column (Phenomenex) and a 10-100 % gradient elution of acetonitrile and 0.1% TFA in water over 45 min. Pure fractions were pooled, lyophilized, and stored at -20 ºC until further use. Identity and purity of peptides were confirmed using analytical HPLC and mass spectrometry under negative ion mode (Agilent 6545 LC/Q-TOF). The mass of the purified peptide was determined, and 5-20 mM stocks were made in DMSO (low enough concentration to assure lipopeptide is fully dissolved). These stocks were aliquoted and stored at -20 ºC.

*Direct Binding Assays*

Direct binding assays were performed using fluorescence polarization in technical triplicate as previously described.[2, 6] Data was analyzed using GraphPad Prism 9.0. For direct binding experiments, a binding curve that accounts for ligand depletion was fit to the observed polarization values as a function of the concentration of protein to obtain the apparent dissociation constant, Kd:

$$y = c + (b - c) \times \frac{(Kd\ +\ a\ +\ x)\ -\ \sqrt{(Kd\ +\ a\ +\ x)^2 - 4ax}}{2a}$$

”y” is the observed total polarization at a given protein concentration, “c” is the minimum observed total polarization value, “b” is the maximum observed total polarization value, “a” is the total concentration of fluorescently labeled peptide, and “x” is the total concentration of protein.

*Competition Binding Assays*

Competition binding assays were performed using fluorescence polarization in technical triplicate as previously described with slight modifications.[6] Each experiment used 20 nM of FITC-labelled peptide and a concentration of protein equivalent to either 1x or 3x the $K_d$ (dependent on protein) were precomplexed and incubated in the presence of serially diluted lipopeptide (50 nM to 400 μM). Total polarization was measured as stated above and fit to a non-linear regression using the built-in equation "log(inhibitor) vs. response – variable slope (four parameters)" (shown below) with the bottom constraint set to 'constant or equal to 35' (as defined in Pherastar parameters) from which the $IC_{50}$ values were calculated. The $IC_{50}$ values were converted to $K_i$ values using the apparent Kd value calculated from the direct binding experiments of the specific coactivator•activator PPIs using a $K_i$ calculator.[7] Reported $K_i$ values are the average of three biological with the indicated error representing the standard deviation of the triplicates.

$$y = Bottom + (Top - Bottom)/(1 + 10^{(LogIC50 - X)*HillSlope})$$

*Differential Scanning Fluorimetry*

Experiments were performed in technical triplicate of 20 μL sample volumes in a 96 well PCR plates sealed with clear cap strips.[8] Assay buffer used was 10 mM sodium phosphate, 100 mM sodium chloride, 10% glycerol, 0.001% NP40, and pH 6.8. To determine $T_m$, 8 μM protein in the presence of 5X SYPRO orange dye (1:1000 dilution in buffer of purchased 5000X stock in DMSO; Invitrogen) was incubated alone or with ligands 34913-8 and 34913-9 (2, 3, 5, 7, 10X) at RT for 30 minutes. An Applied Biosystems StepOnePlus qPCR instrument was used to obtain melting curves by excitation at 488 nm and emission measured at 602 nm over a temperature gradient of 25–95°C with a 1 °C/min increase. Raw fluorescence data was imported into the online data analysis program, DSFworld, and $T_m$ was calculated by determining the maximum of the first derivative (dRFU).[9],[10] For data visualization, raw fluorescence units and dRFU was plotted as a function of temperature using GraphPad Prism software. Change in melting temperature ($\Delta T_m$) of each ligand was calculated as the difference between the $T_m$ of the protein and the $T_m$ of the protein + ligand. Reported values are the averages of biological duplicates and their indicated error representing the standard deviation of the duplicates.

*NMR Spectroscopy*

NMR assignments of Med25 AcID (395-545) were determined from previous studies using $^{13}C$, $^{15}N$-labeled protein.[2, 11] Constant time $^{1}H,^{15}N$-HSQC and $^{1}H,^{13}C$-HSQC experiments were performed using 75 μM $^{13}C,^{15}N$-labeled Med25 in NMR buffer (20 mM sodium phosphate pH 6.5, 150 mM NaCl, 3 mM DTT, 10% D2O, and 2% DMSO) on a Bruker 600 MHz instrument equipped with a cryoprobe. Data processing and visualization was performed using NMRPipe[12] and NMRFAM-Sparky,[13] respectively. Peak assignments of lipopeptide analog-bound complexes were achieved by titration experiments with the lead lipopeptidomimetics (34913-8 and 34913-9) with titration points of 0.2, 0.5, 0.8, 1.1, 2, and 3 equivalents of compound. Control spectra were obtained with Med25 and DMSO only. Chemical shift perturbations ($\Delta_\delta$) were calculated from the proton ($\Delta_\delta H$) and carbon ($\Delta_\delta C$) chemical shifts by:

$$\Delta\delta = \sqrt{(\Delta\delta_H)^2 + (0.25 \times \Delta\delta_C)^2}$$

*Mammalian Cell Culture*

VARI068 cells are from a culture of a triple negative breast cancer patient-derived xenograft as previously described.[14],[15] VARI068 cells were grown in Dulbecco's Modified Eagle Medium (Gibco, cat. #: 11965092) supplemented with 10% fetal bovine serum (FBS), 1x Antibiotic-Antimycotic and 10mg/mL gentamicin in a 37 ºC incubator with 5% carbon dioxide.

*Cellular Thermal Shift Assays*

VARI068 cells were harvested using standard protocols and pelleted (~4 million cells per pellet) in 1.5 mL microcentrifuge tubes at 1500 x g for 5 minutes at 4 ºC. Nuclear extracts were generated according to the manufacturer's protocols using NE-PER Nuclear and Cytoplasmic Extraction Reagents (Thermo Fisher Scientific, cat #: 78833). Nuclear extracts were buffer exchanged into phosphate buffered saline (PBS) using Zeba Desalting Column, 7K MWCO (Thermo Fisher Scientific). Prepared nuclear extracts were evenly split into three samples. 34913-8 and 34913-9 (dissolved in DMSO) were added to two samples at the indicated concentration with equivalent volume of DMSO added to the final sample. Final concentration of DMSO was 0.1% v/v. Dosed nuclear extracts were incubated at room temperature for 30 mins. Following incubation, extracts were aliquoted into thin-walled PCR tubes (20 μL per tube, ~300,000 cells per tube).

A Labnet Multigene OPTIMAX PCR was used to heat samples for 3 minutes at six temperatures: 54 ºC, 58 ºC, 62 ºC, 66 ºC, 70 ºC, 74 ºC. Contents of PCR tubes were centrifuged at 17000 x g for 2 minutes at 4 ºC and transferred, leaving the precipitated protein pellet undisturbed. LDS loading dye was added to supernatant, boiled, loaded onto a 4-20% mini-PROTEAN TGX gel (BioRad), and run at 170V for 1 hour. Protein was transferred from gel to PDVF membrane using standard protocols on a Bio-Rad Trans-Blot Turbo Transfer System. The membrane was blocked for 1 hour at RT with gentle shaking using SuperBlock™ Blocking Buffer in PBS (Thermo Scientific). Super Block was removed and Med25 antibody (Novus Biologicals, NBP2-55868) was added to membrane (1:500 dilution with 1:500 Tween 20 in SuperBlock™) and incubated overnight at 4 ºC with gentle shaking. Primary antibody was removed and the membrane was washed three times with PBST. Secondary antibody (Santa Cruz, sc-2357, 1:1000 with 1:500 Tween 20 in SuperBlock™) was added to the membrane and incubated for 1 hour at RT with gentle shaking. Secondary antibody was removed and membrane was washed three times with PBST. HRP substrate (Thermo Scientific) was added to the membrane and incubated for 1 minute at RT. Western blot was visualized using Chemiluminscence on an Azure Biosystems c600 imager. Blots were analyzed using ImageJ software. Reported values are representative of one biological duplicate, with a second biological duplicate found in SI (Figure S20).

*Quantitative Polymerase Chain Reaction*

For endogenous gene expression analysis, VARI068 cells were seeded in a 24-well plate ($1x10^5$ cells/well) and allowed to adhere overnight at 37 ºC. Media was removed and replaced with fresh media containing vehicle or compound delivered in DMSO (0.2% v/v) at the indicated concentrations. After 3 hours, the media was removed, and total RNA was isolated using RNeasy Plus Mini Kits (Qiagen) following the manufacturer's protocol. Each RNA sample was used to synthesize cDNA using iScript Reverse Transcription Supermix (Bio-Rad). RT-qPCR samples were run in technical triplicate on an Applied Biosystems StepPlusOne instrument using the cDNA, GoTaq qPCR Master Mix (Promega).

| **Primers** | **Sequence (5'-3')** |
| --- | --- |
| *Human RPL19 Forward* | ATGTATCACAGCCTGTACCTG |
| *Human RPL19 Reverse* | TTCTTGGTCTCTCTTCCTCCTTG |

| | |
|---|---|
| *MMP2 Forward* | CATTCCAGGCATCTGCGATGAG |
| *MMP2 Reverse* | AGCGAGTGGATGCCGCCTTTAA |

*Table S2.* Gene primers used for qPCR and their sequences.

RT-PCR analysis was performed using the comparative CT method (ΔΔCT Method) to estimate MMP2 mRNA transcript levels compared to control RPL19 mRNA transcript levels. Reported values are the averages of biological triplicates and their indicated error representing the standard deviation of the triplicates.

**SI Figures**

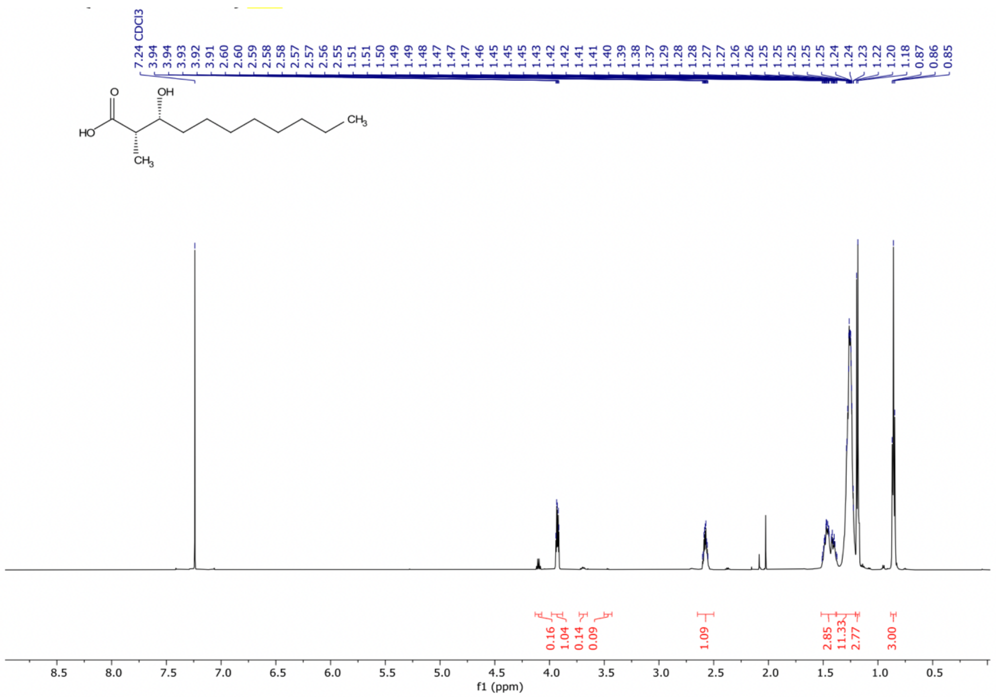


*Figure S1.* $^{1}H$ NMR (600 MHz, $CDCl_3$) of **S2**.

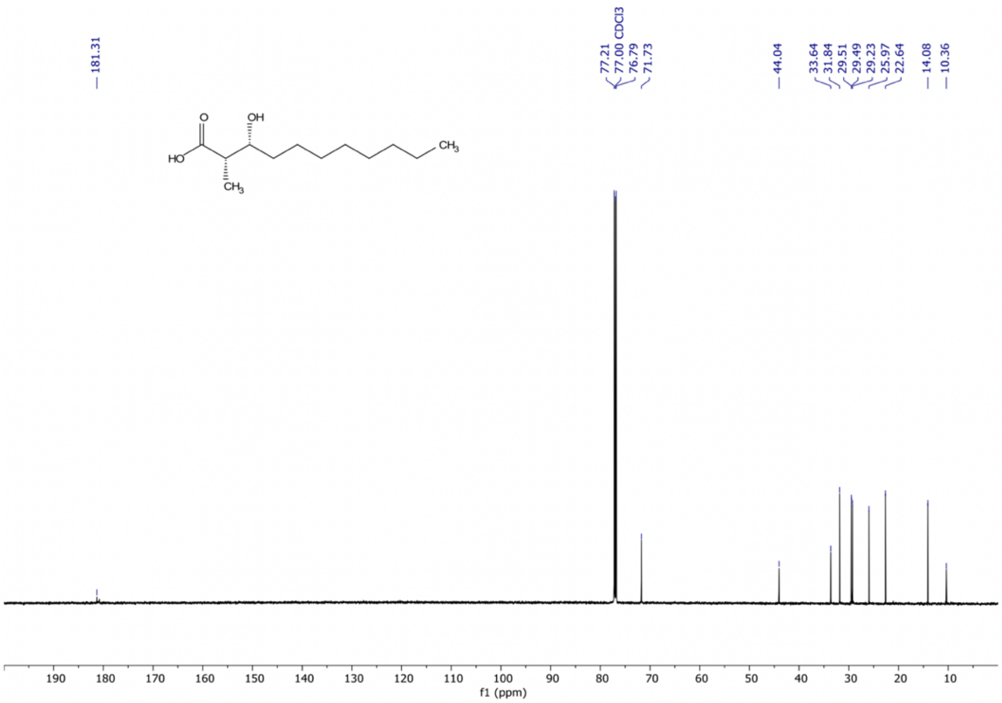


*Figure S2.* $^{13}C$ NMR (15 MHz, $CDCl_3$) of **S2**.

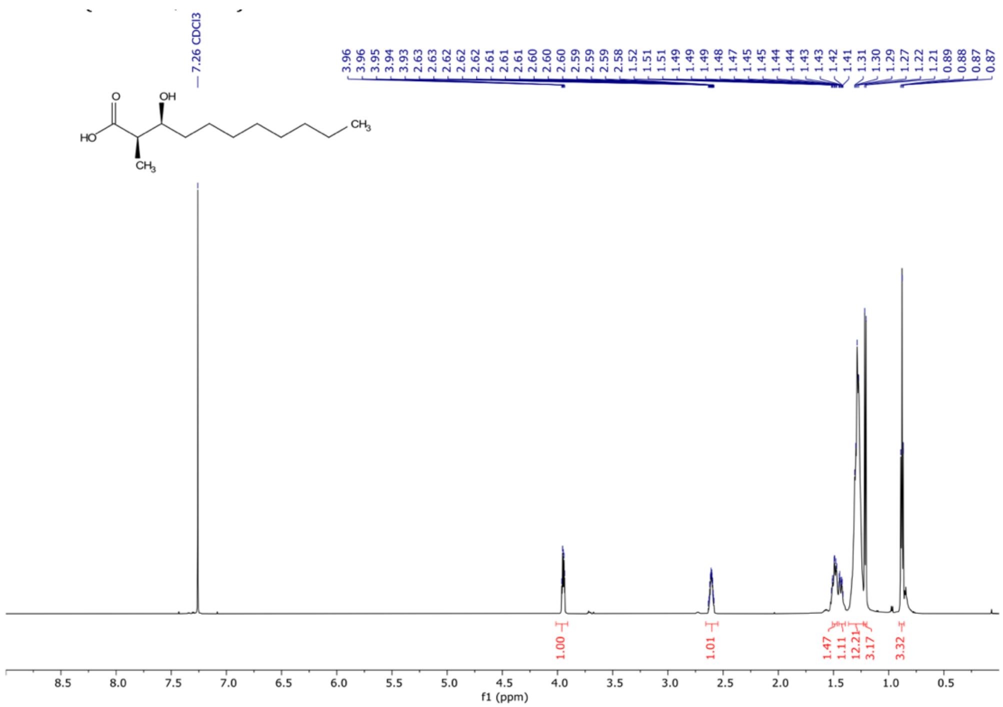

*Figure S3.* $^{1}H$ NMR (600 MHz, $CDCl_3$) of **S4**.

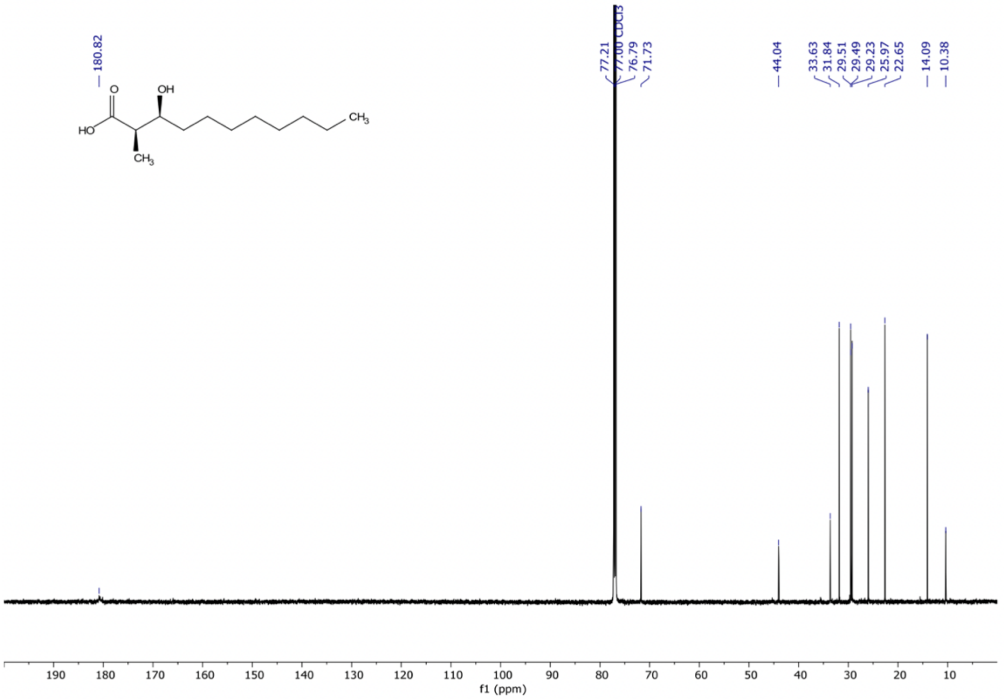

*Figure S4.* $^{13}C$ NMR (150 MHz, $CDCl_3$) of **S4**.

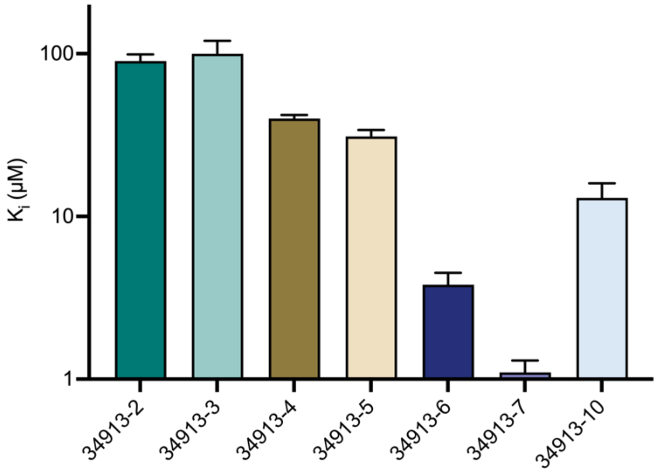


*Figure S5.* Inhibition of Med25 AcID•ATF6α by lipopeptidomimetics as determined by competitive fluorescence polarization assays. Apparent $IC_{50}$ values were determined through titration of compound for Med25 AcID•ATF6α in experimental triplicate with the indicated error (SD). The $IC_{50}$ values were converted to $K_i$ values using the apparent $K_d$ value based on the direct binding of Med25 AcID•ATF6α.[7] Data shown is the average of three independent experiments with the indicated error (SD).

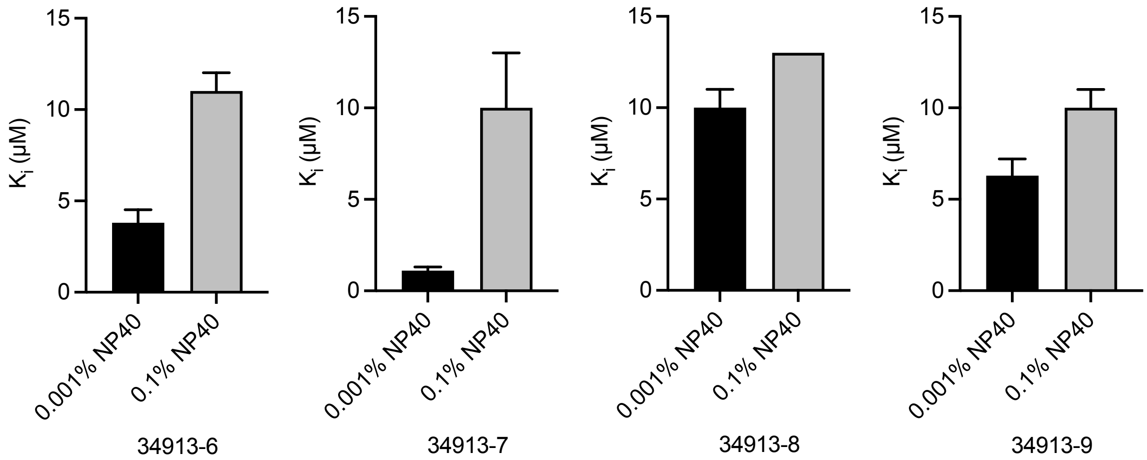


*Figure S6.* Inhibition of Med25 AcID•ATF6α by 34913-6 (left), 34913-7 (left middle), 34913-8 (right middle), and 34913-9 (right) with different NP40 concentrations as determined by competitive fluorescence polarization assays. Apparent $IC_{50}$ values were determined through titration of compound for Med25 AcID•ATF6α PPI performed in experimental triplicate with the indicated error (SD). The $IC_{50}$ values were converted to $K_i$ values using the apparent $K_d$ value based on the direct binding of Med25 AcID•ATF6α PPI using a $K_i$ calculator.[7] Data shown is the average of two independent experiments with the indicated error (SD).

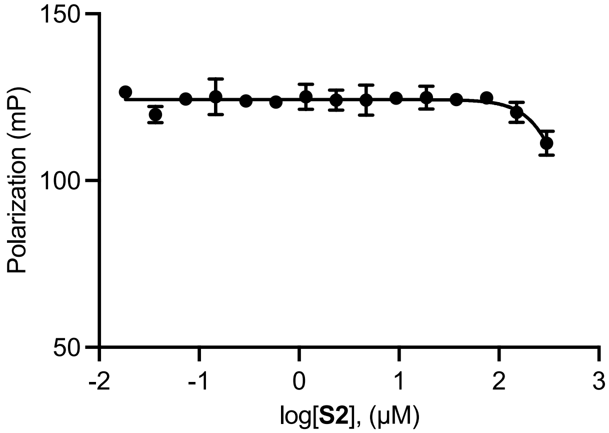


*Figure S7.* Inhibition of Med25 AcID•ATF6α by **S2** as determined by competitive fluorescence polarization assays. Curve represents the mean value of three technical triplicates with vertical error bars representing the standard deviation of the fraction of tracer bound at the indicated **S2** concentration.

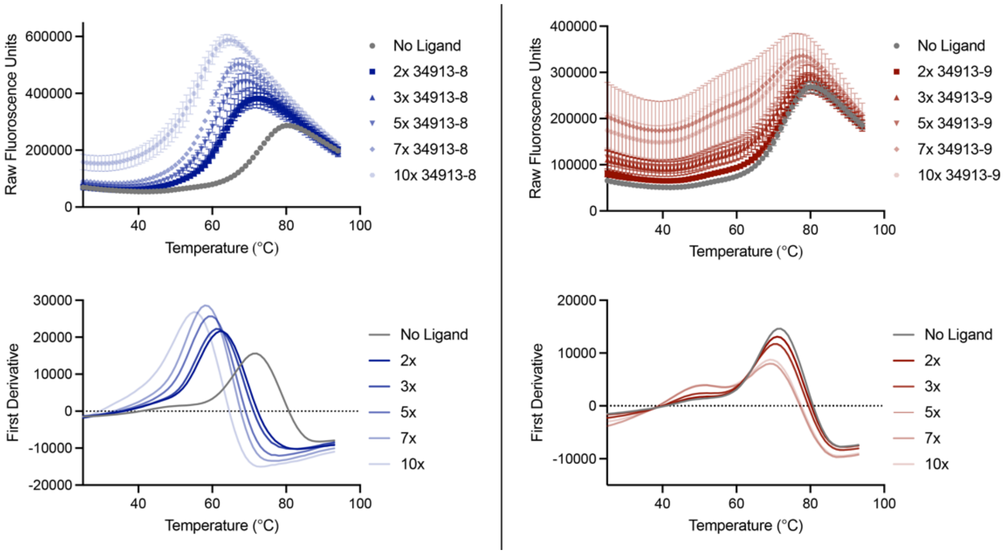


*Figure S8.* Raw DSF data for lead lipopeptidomimetics, 34913-8 and 34913-9. Left. Raw fluorescence units (top) and first derivative (bottom) traces of Med25 AcID 2, 3, 5, 7, and 10 equiv. of 34913-8. Right. Raw fluorescence units (top) and first derivative (bottom) traces of Med25 AcID 2, 3, 5, 7, and 10 equiv. of 34913-9. Melting temperature was determined from raw fluorescence data using DSFworld.[9-10] Data was obtained in technical triplicates with the indicated error (SD) and is representative of experiments performed in biological duplicates.

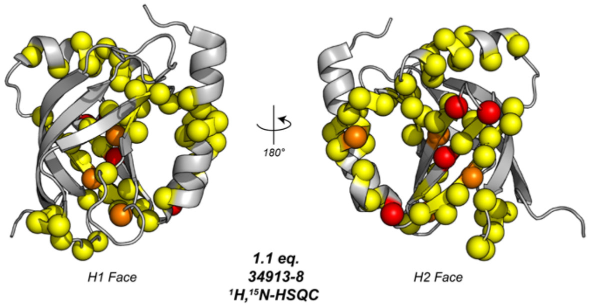


*Figure S9.* All $^{1}$H, $^{15}$N-HSQC perturbations of 1.1 equiv. of 34913-8 mapped onto Med25 AcID (PDB ID 2XNF). All residues with CSPs above the signal-to-noise (S/N) ratio (≥ 0.02 ppm) are shown, including several residues with weaker shift patterns (yellow residues). Yellow = 0.02 ppm – 0.085 ppm, orange = 0.0851 ppm – 0.14 ppm, red ≥ 0.141 ppm.

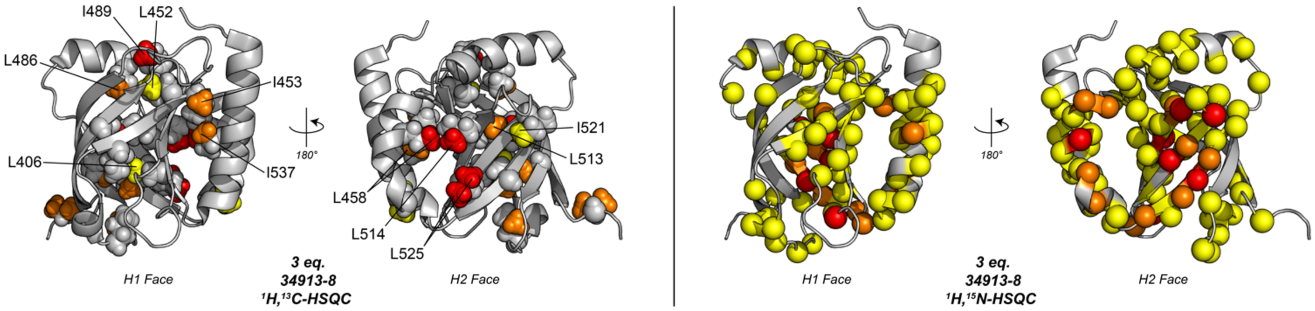


*Figure S10***.** Chemical shifts perturbations from saturated concentrations of 34913-8 (3 equiv.) mapped onto Med25 AcID (PDB ID 2XNF). Left. $^{1}$H,$^{13}$C-HSQC spectra with high concentrations of 34913-8 showed increased chemical shift perturbations, though significant perturbations (orange and red residues) occurred primarily on the H2 face of AcID and flanking α-helices. Yellow = 0.02 ppm - 0.0249 ppm, orange = 0.025 ppm - 0.049 ppm, red ≥ 0.0.0491 ppm. Right. $^{1}$H, $^{15}$N-HSQC spectra revealed several significant changes to the H2 face, with only minor shifts affecting the H1 face (yellow residues). Yellow = 0.02 ppm - 0.085 ppm, orange = 0.0851 ppm - 0.14 ppm, red ≥ 0.141 ppm.

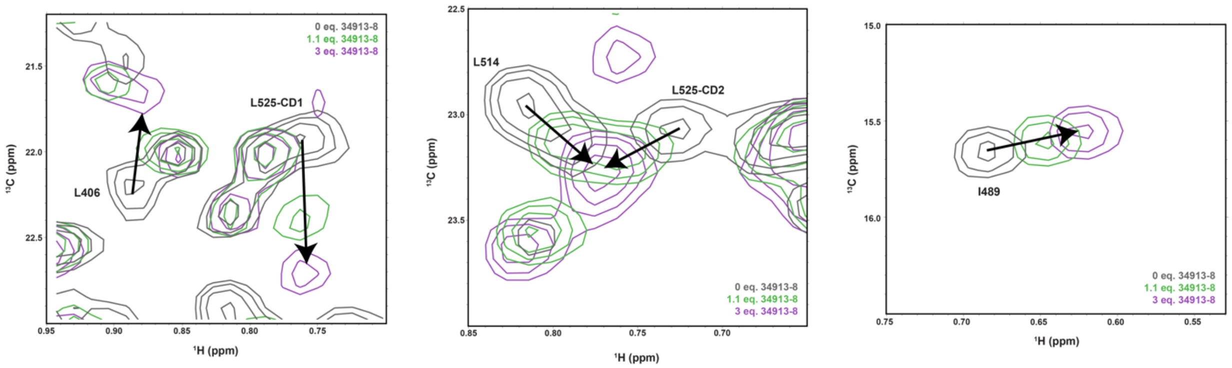


*Figure S11.* Overlay of $^{1}$H, $^{13}$C-HSQC spectra of free Med25 (dark grey), 1.1 equiv. of 34913-8 (green), 3 equiv. of 34913-8 (purple). Peaks that have significant CSPs at variable equiv. of compound are labelled.

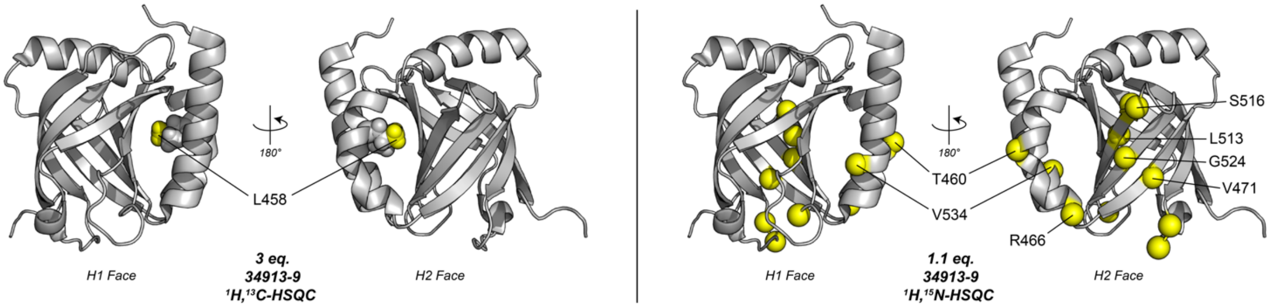


*Figure S12***.** $^{1}$H, $^{13}$C and $^{1}$H, $^{15}$N-HSQC CSPs induced by binding of 34913-9 mapped onto Med25 AcID (PDB ID 2XNF). $^{1}$H, $^{13}$C and $^{1}$H, $^{15}$N-HSQC experiments show that 34913-9 does not significantly interact with Med25 AcID. Left. $^{1}$H, $^{13}$C CSPs plotted onto Med25 AcID (PDB ID 2XNF) reveal one residue (L458) with a notable CSP at 3 equiv. of lipopeptide. Yellow = 0.02 ppm - 0.0249 ppm, orange = 0.025 ppm - 0.049 ppm, red ≥ 0.0.0491 ppm. Right. $^{1}$H, $^{15}$N CSPs indicate minor perturbations primarily on the H2 face β-barrel and flanking α-helices at 1.1 equiv. of 34913-9. Yellow = 0.02 ppm - 0.085 ppm, orange = 0.0851 ppm - 0.14 ppm, red ≥ 0.141 ppm.

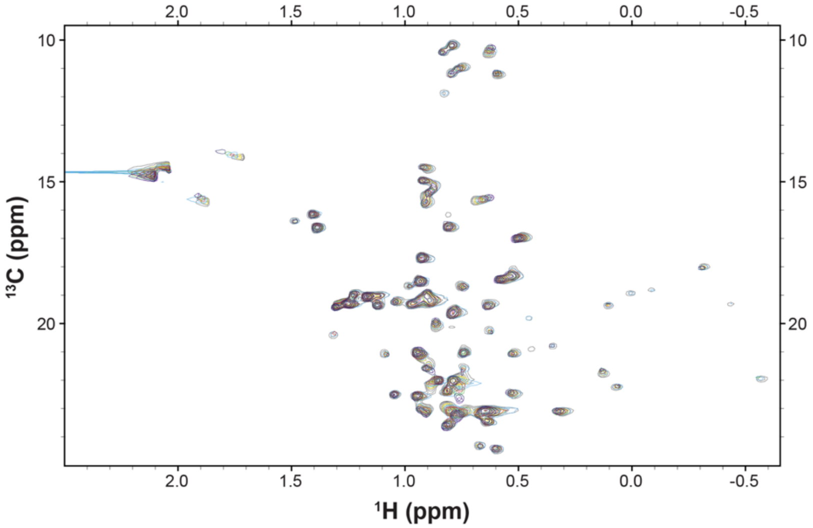


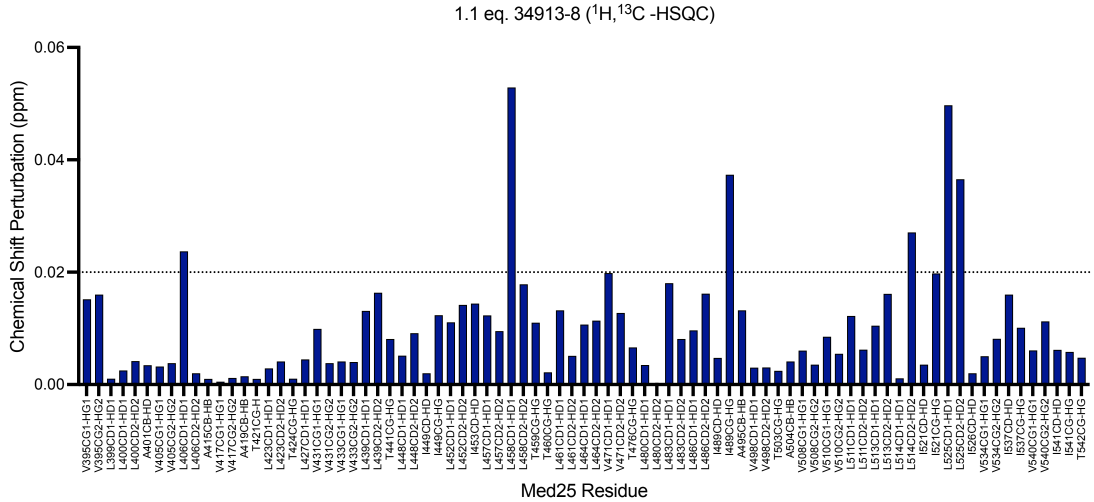


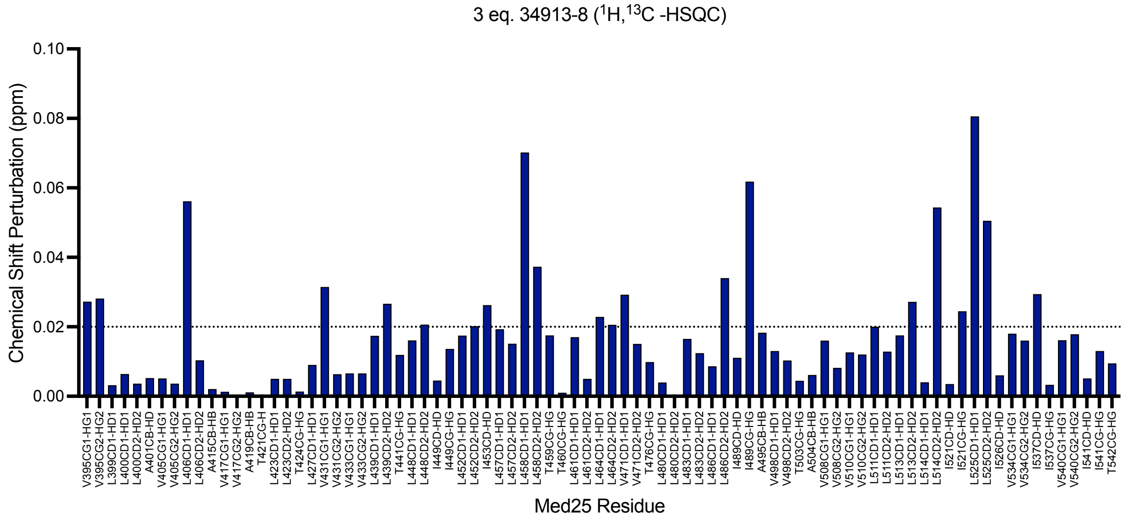


*Figure S13.* $^{1}$H, $^{13}$C-HSQC spectra of 34913-8 and all residues CSPs. Top. Overlay of $^{1}$H, $^{13}$C-HSQC spectra of Med25 with titration of 34913-8. Spectra shown are free Med25 (dark grey), 0.2 equiv. of 34913-8 (yellow), 0.5 equiv. 34913-8 (light blue), 0.8 equiv. 34913-8 (red), 1.1 equiv. 34913-8 (green), 2 equiv. 34913-8 (dark blue), 3 equiv.

34913-8 (purple). Middle. $^{1}$H, $^{13}$C-HSQC CSPs of Med25 residues with 1.1 equiv. of 34913-8. Bottom. $^{1}$H, $^{13}$C-HSQC CSPs of Med25 residues with 3 equiv. of 34913-8. Dashed line indicates the level of 3 standard deviations above the average chemical shift perturbation.

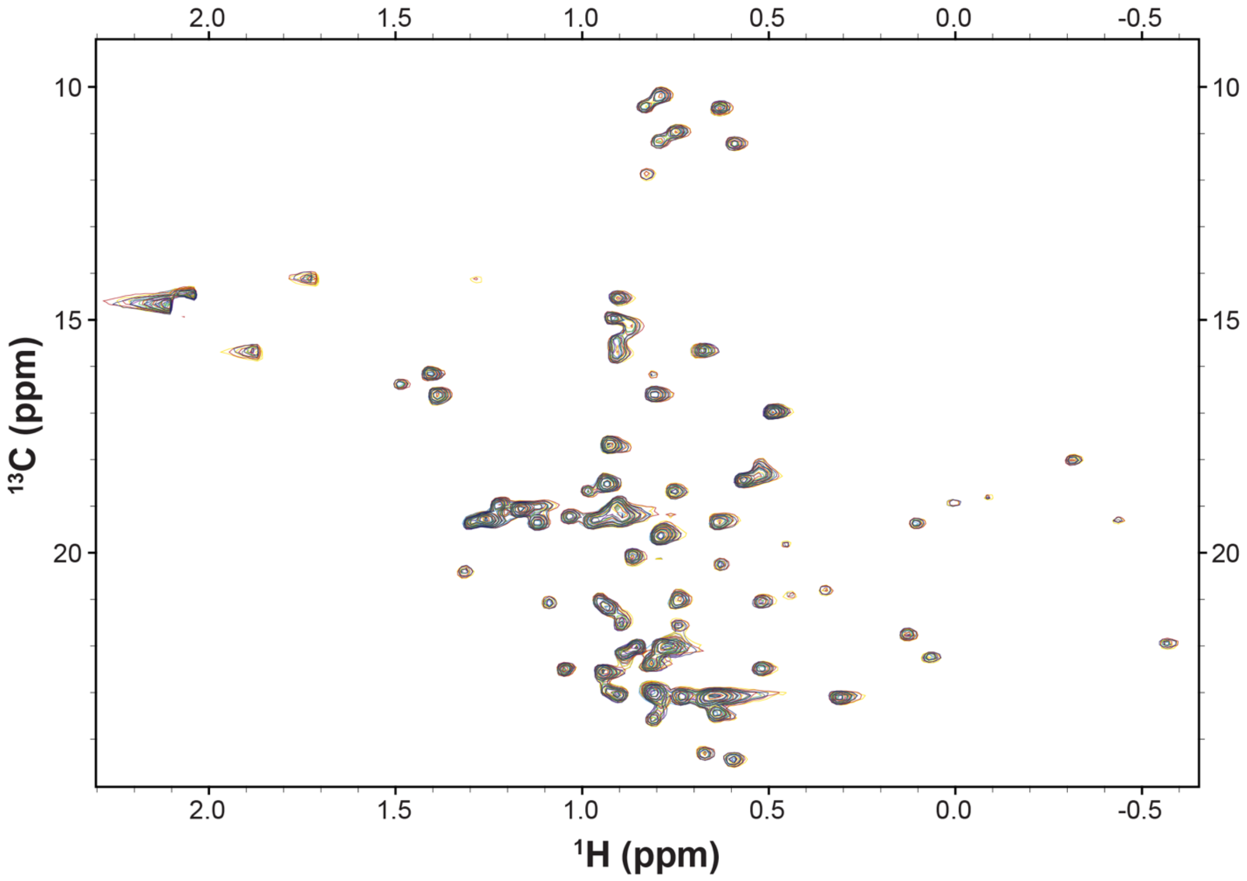


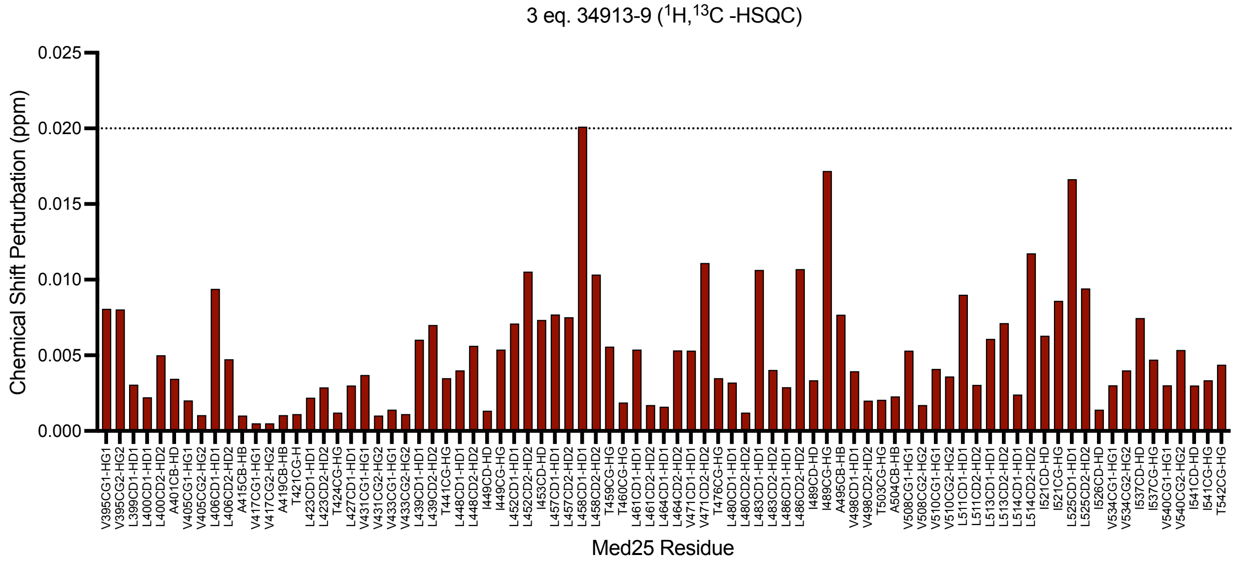


*Figure S14.* $^{1}$H, $^{13}$C-HSQC spectra of 34913-9 and all residue CSPs. Top. Overlay of $^{1}$H, $^{13}$C-HSQC spectra of Med25 with titration of 34913-9. Spectra shown are free Med25 (dark grey), 0.2 equiv. of 34913-9 (yellow), 0.5 equiv. 34913-9 (light blue), 0.8 equiv. 34913-9 (red), 1.1 equiv. 34913-9 (green), 2 equiv. 34913-9 (dark blue), 3 equiv. 34913-9 (purple). Bottom. $^1$H, $^{13}$C-HSQC CSPs of Med25 residues with 3 equiv. of 34913-9. Dashed line indicates the level of 3 standard deviations above the average chemical shift perturbation.

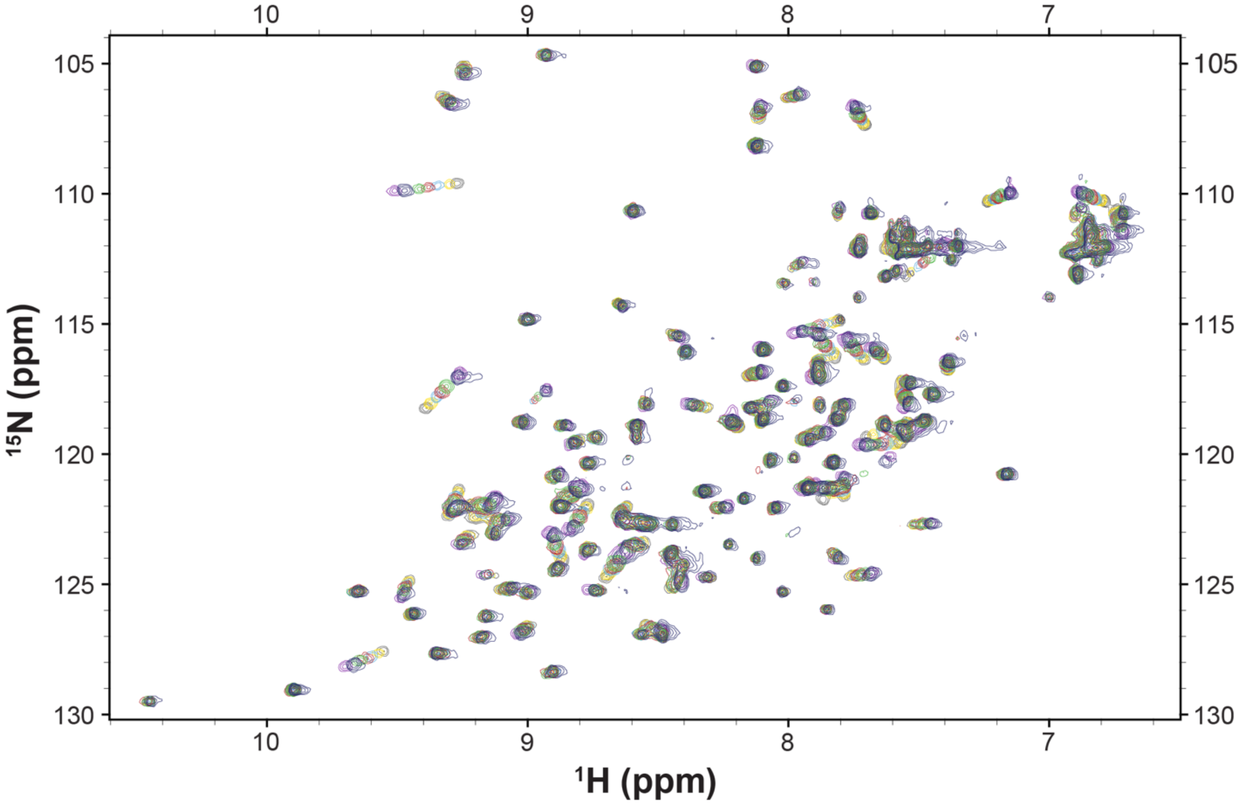


*Figure S15.* Overlay of $^1$H, $^{15}$N-HSQC spectra of Med25 with titration of 34913-8. Spectra shown are free Med25 (dark grey), 0.2 equiv. of 34913-8 (yellow), 0.5 equiv. 34913-8 (light blue), 0.8 equiv. 34913-8 (red), 1.1 equiv. 34913-8 (green), 2 equiv. 34913-8 (dark blue), 3 equiv. 34913-8 (purple).

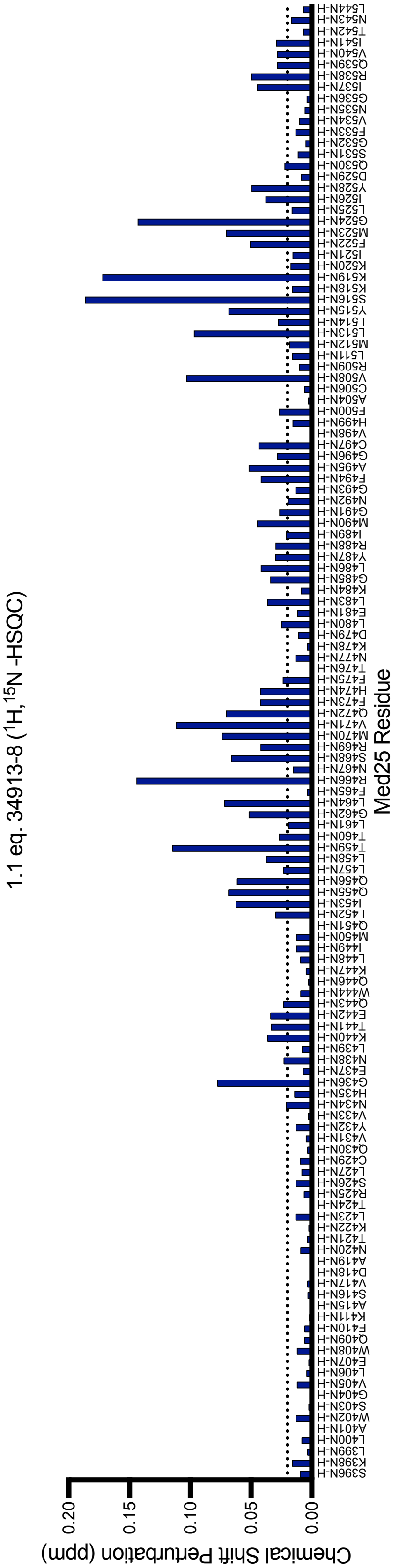


*Figure S16* $^{1}$H,$^{15}$N-HSQC CSPs of Med25 residues with 1.1 equiv. of 34913-8. Dashed line indicates the level of 3 standard deviations above the average chemical shift perturbation.

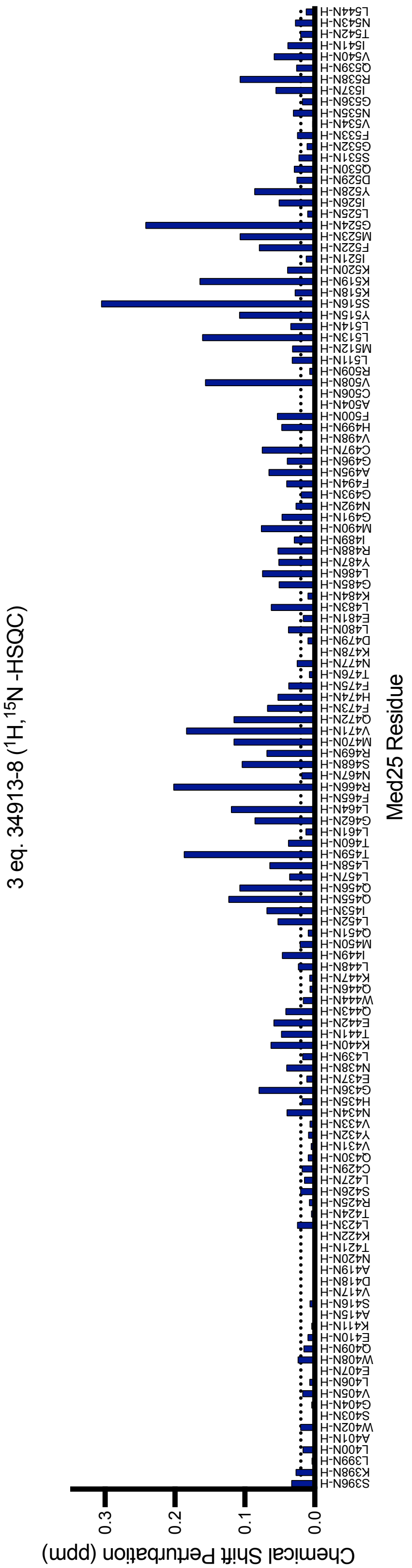


*Figure S17.* $^{1}$H, $^{15}$N-HSQC CSPs of Med25 residues with 3 equiv. of 34913-8. Dashed line indicates the level of 3 standard deviations above the average chemical shift perturbation.

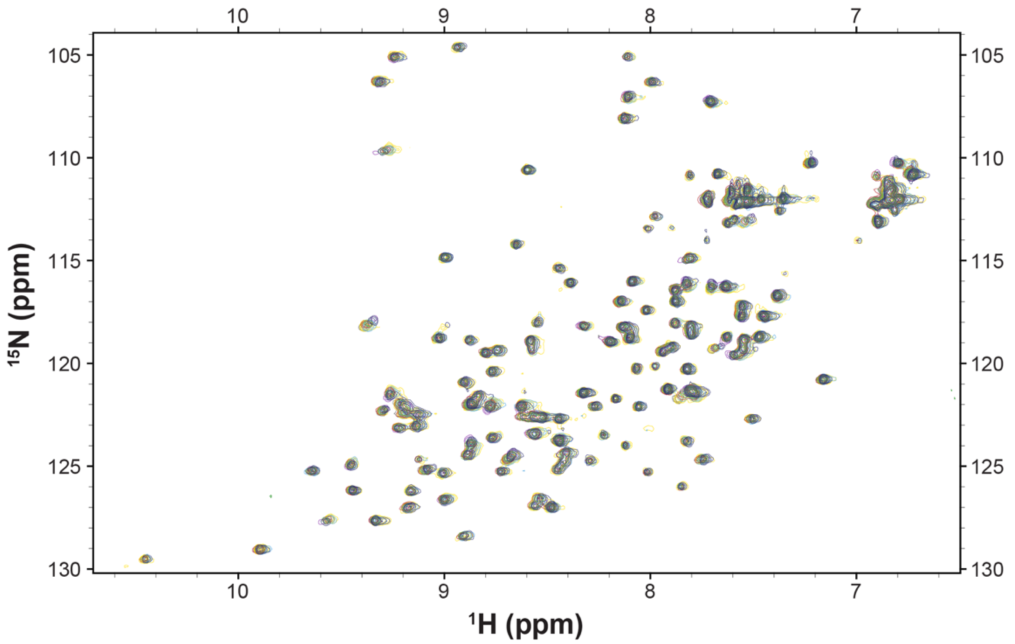


*Figure S18.* Overlay of $^{1}$H, $^{15}$N-HSQC spectra of Med25 with titration of 34913-9. Spectra shown are free Med25 (dark grey), 0.2 equiv. of 34913-9 (yellow), 0.5 equiv. 34913-9 (light blue), 0.8 equiv. 34913-9 (red), 1.1 equiv. 34913-9 (green), 2 equiv. 34913-9 (dark blue), 3 equiv. 34913-9 (purple).

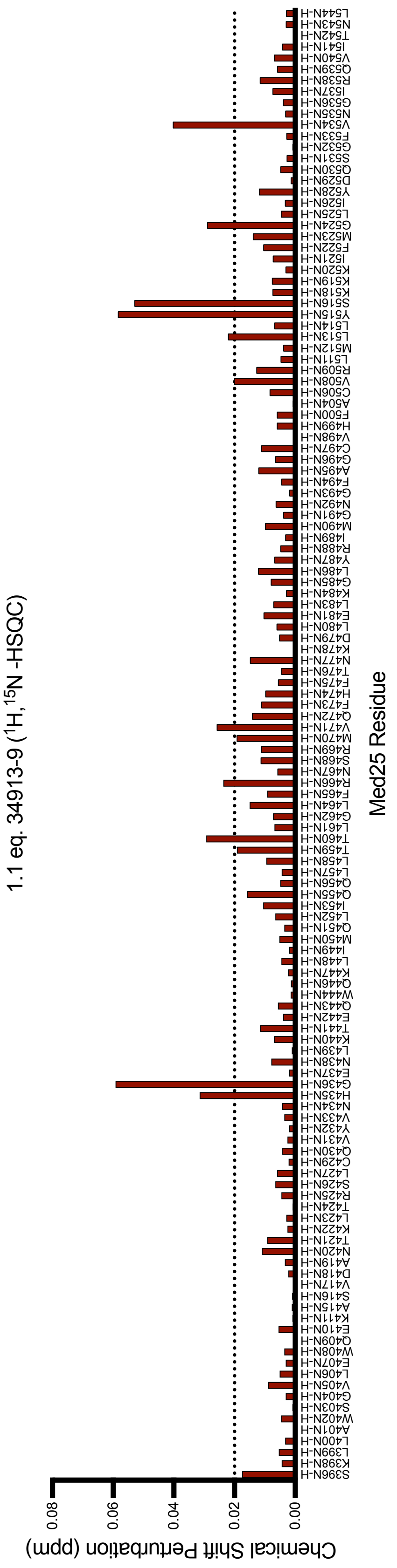


*Figure S19.* [1]H, [15]N-HSQC CSPs of Med25 residues with 1.1 equiv. of 34913-9. Dashed line indicates the level of 3 standard deviations above the average chemical shift perturbation.

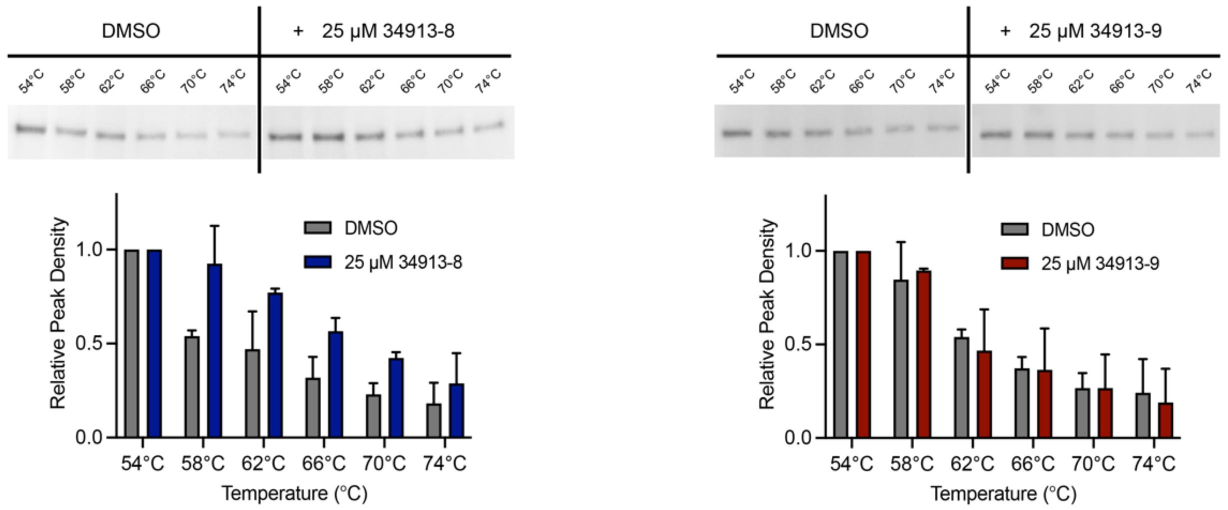


*Figure S20.* Cellular thermal shift assays of 34913-8 and 34913-9 using VARI068 nuclear extracts. Experiments were conducted at 25 µM of lipopeptidomimetic compared to DMSO control (biological duplicates). Western blots show thermal stabilization for Med25 with 34913-8 (top left) and not 34913-9 (top right). Band densities, calculated using ImageJ, demonstrate 2-fold stability with 34913-8 (bottom left) while 34913-9 shows similar band densities to DMSO control (bottom right). Error bars represent the standard deviation of the mean from biological duplicates.

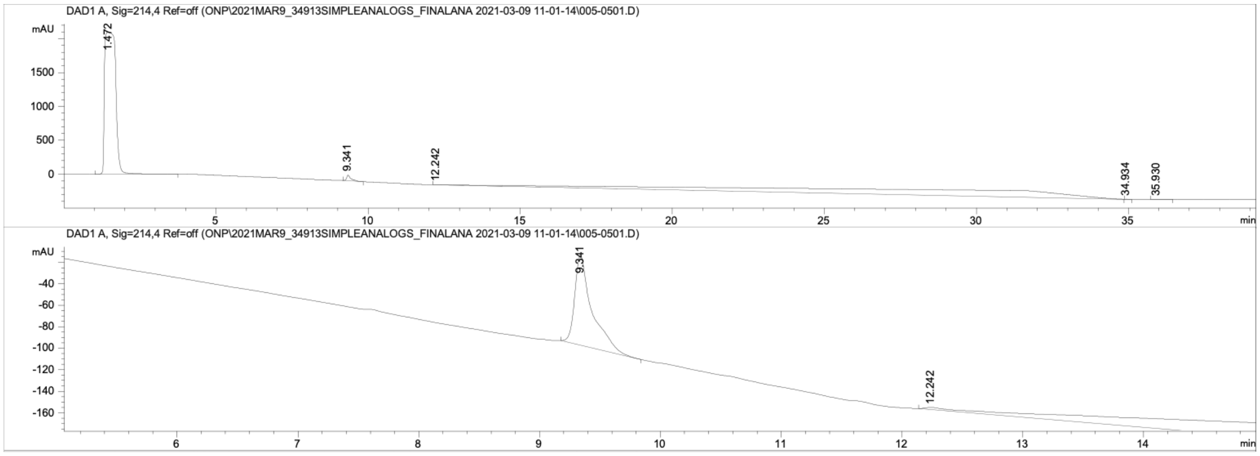


*Figure S21.* Top. Analytical trace of **34913-2** at > 97% purity at 214 nm. Bottom. Zoomed in analytical trace at 214 nm. Peak before 5 min. corresponds to solvent peak, DMSO, that was used to dissolve sample.

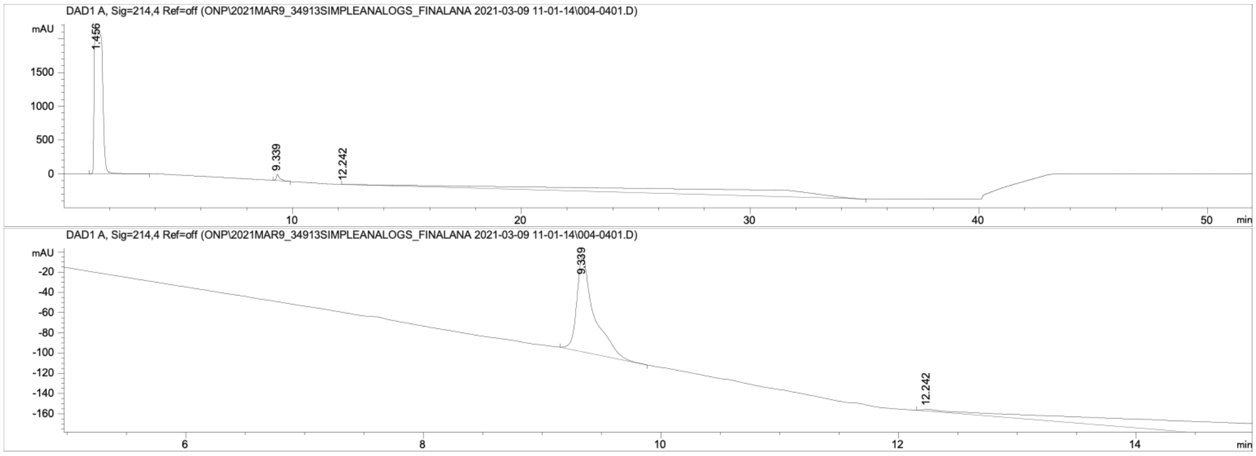


*Figure S22.* Top. Analytical trace of **34913-3** at > 97% purity at 214 nm. Bottom. Zoomed in analytical trace at 214 nm. Peak before 5 min. corresponds to solvent peak, DMSO, that was used to dissolve sample.

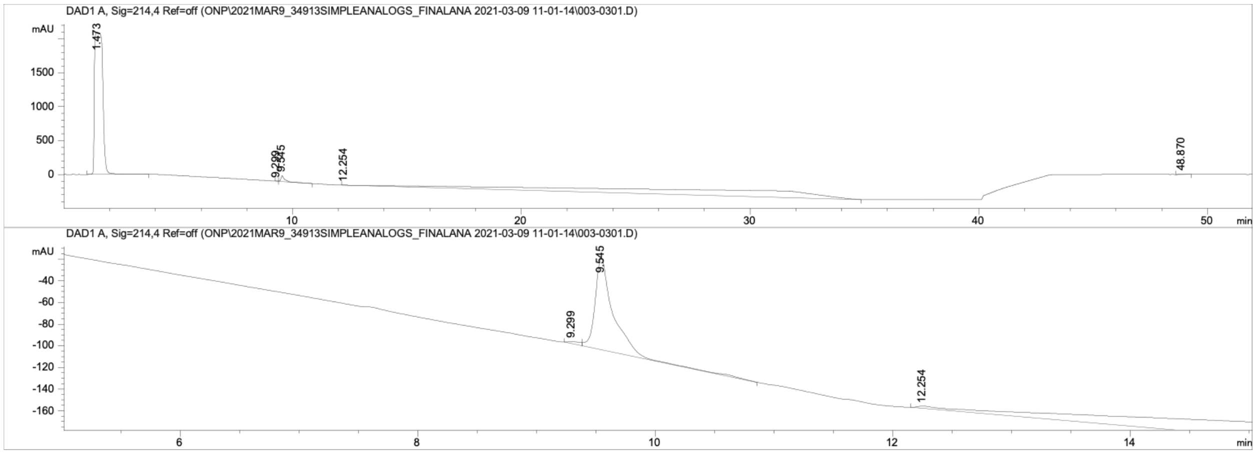

*Figure S23.* Top. Analytical trace of **34913-4** at > 97% purity at 214 nm. Bottom. Zoomed in analytical trace at 214 nm. Peak before 5 min. corresponds to solvent peak, DMSO, that was used to dissolve sample.

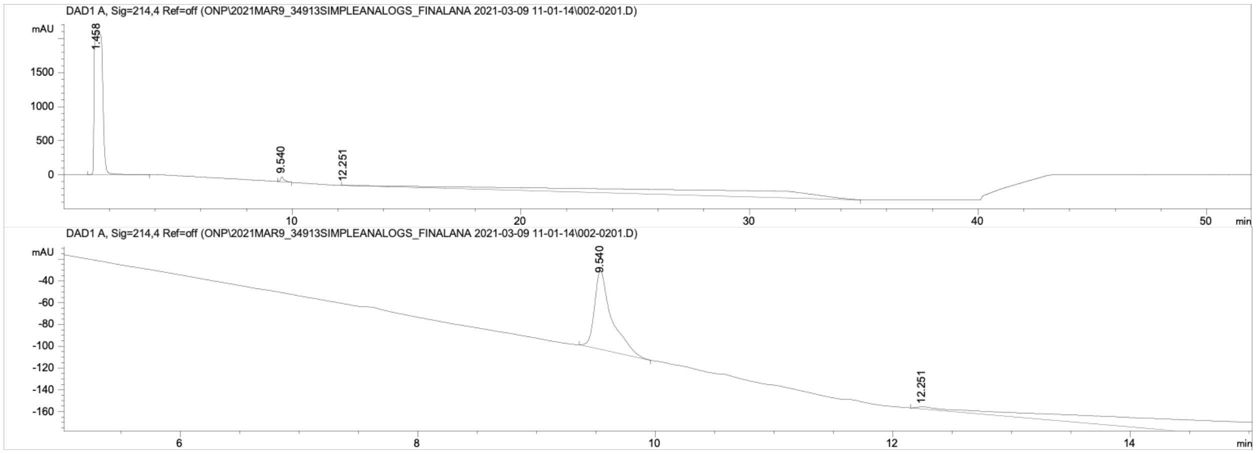


*Figure S24.* Top. Analytical trace of **34913-5** at >97% purity at 214 nm. Bottom. Zoomed in analytical trace at 214 nm. Peak before 5 min. corresponds to solvent peak, DMSO, that was used to dissolve sample.

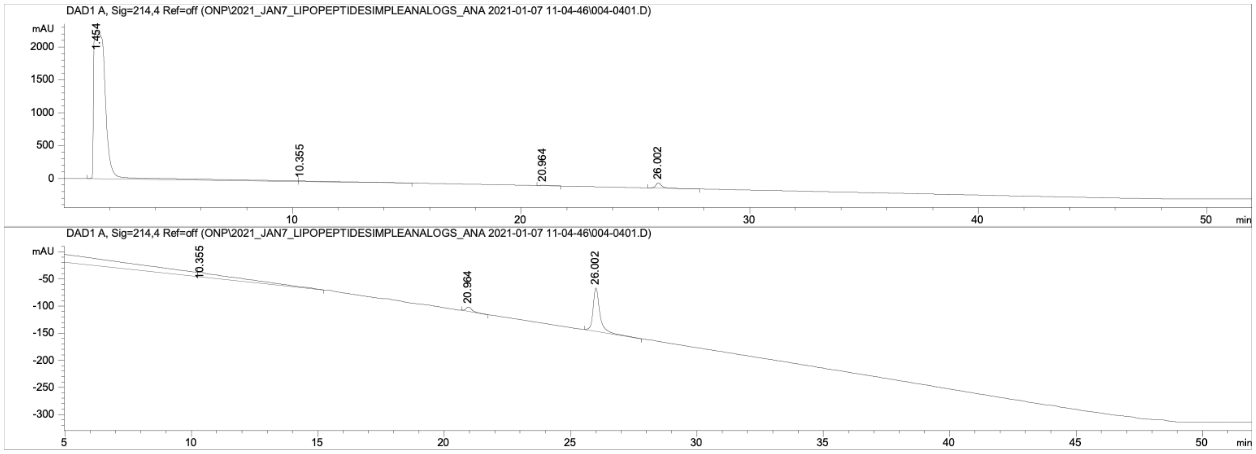


*Figure S25.* Top. Analytical trace of **34913-6** at > 90% purity at 214 nm. Bottom. Zoomed in analytical trace at 214 nm. Peak before 5 min. corresponds to solvent peak, DMSO, that was used to dissolve sample.

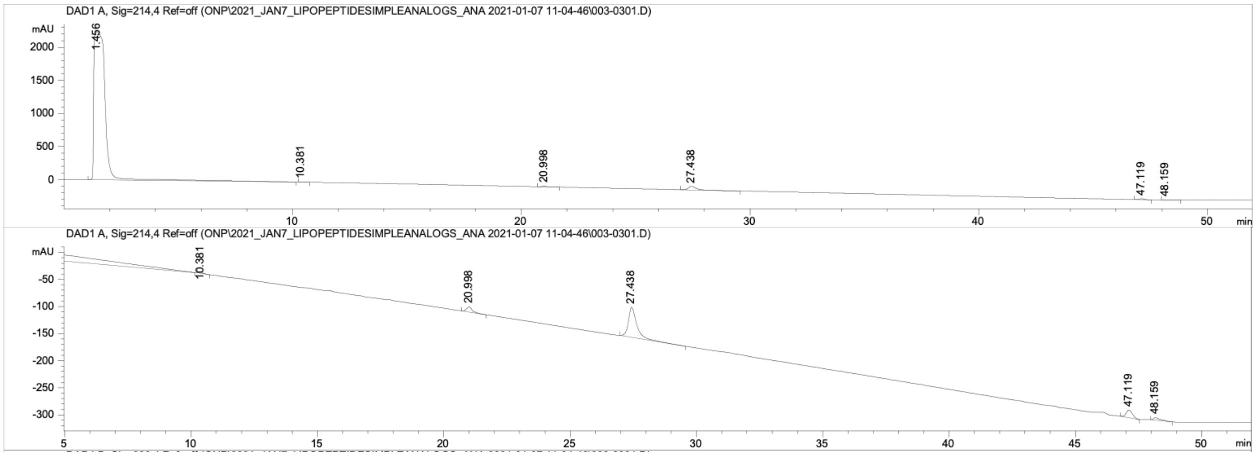

*Figure S26.* Top. Analytical trace of **34913-7** at > 90% purity at 214 nm. Bottom. Zoomed in analytical trace at 214 nm. Peak before 5 min. corresponds to solvent peak, DMSO, that was used to dissolve sample.

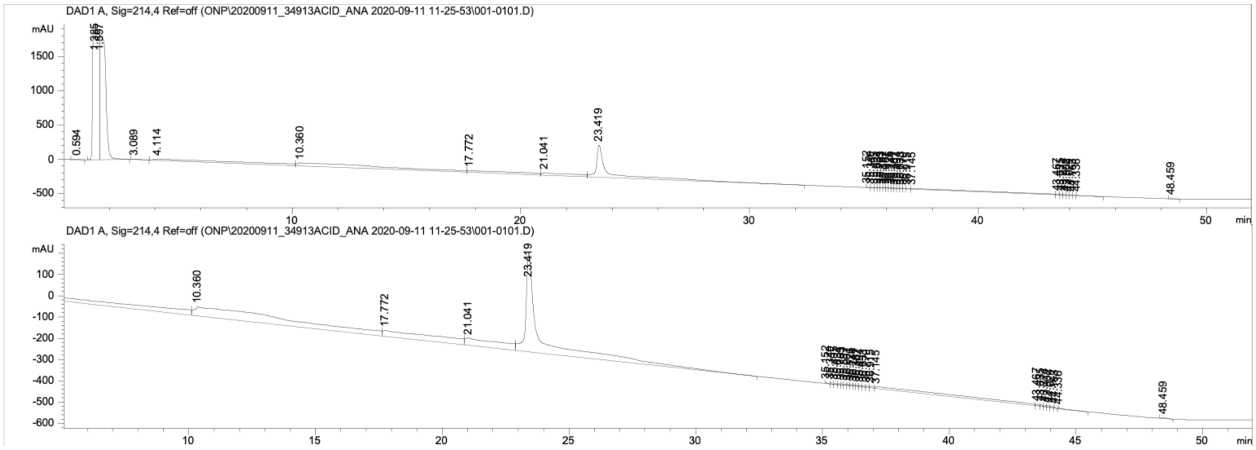

*Figure S28.* Top. Analytical trace of **34913-8** at > 97% purity at 214 nm. Bottom. Zoomed in analytical trace at 214 nm. Peak before 5 min. corresponds to solvent peak, DMSO, that was used to dissolve sample.

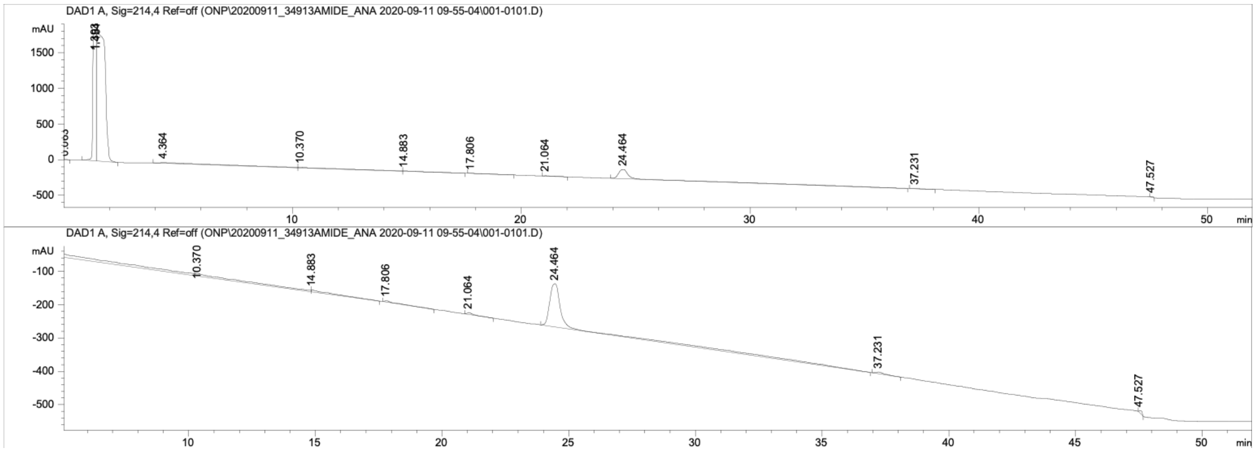


*Figure S29***.** Top. Analytical trace of **34913-9** at > 97% purity at 214 nm. Bottom. Zoomed in analytical trace at 214 nm. Peak before 5 min. corresponds to solvent peak, DMSO, that was used to dissolve sample.

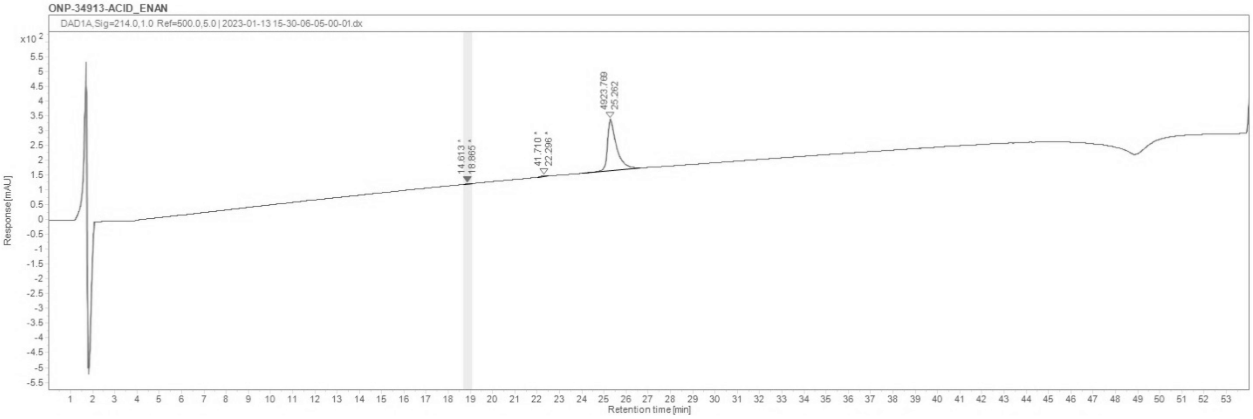


*Figure 30.* Analytical trace of **34913-10** at > 97% purity at 214 nm.

*Analytical Traces of Peptides Used in this Study*

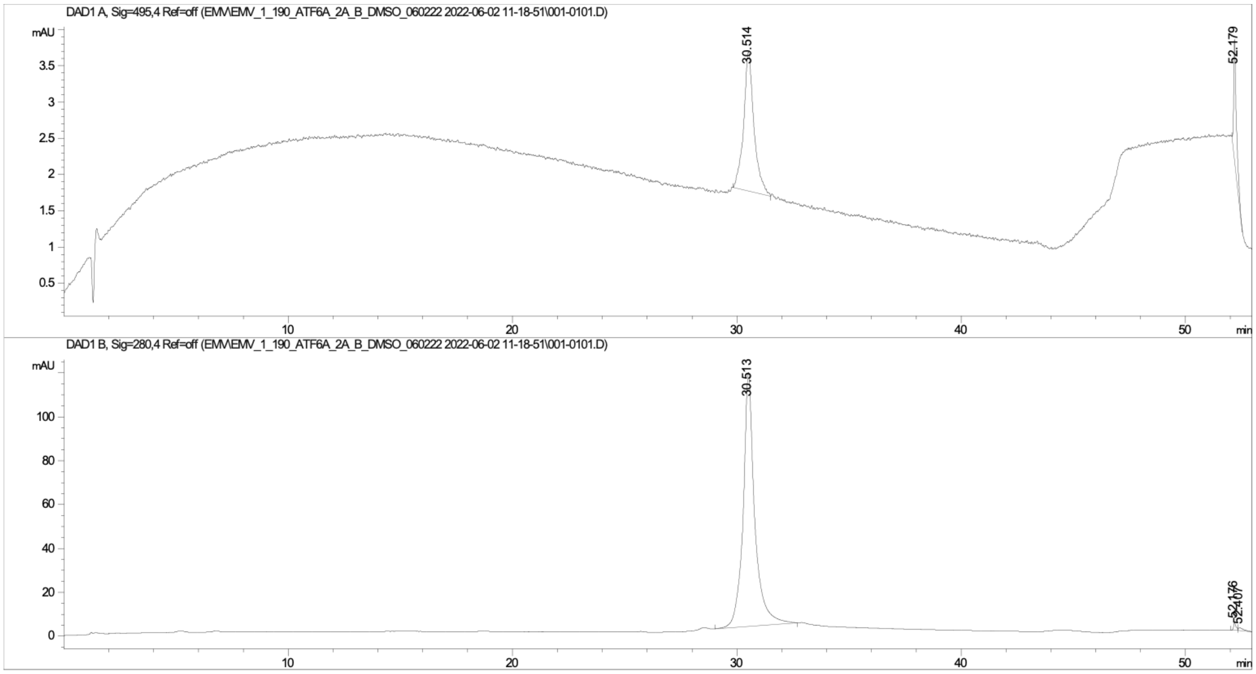


*Figure S31.* Analytical HPLC trace of **FITC-ATF6α (38-75)** monitored at 495 nm (top) and 280 nm (bottom).

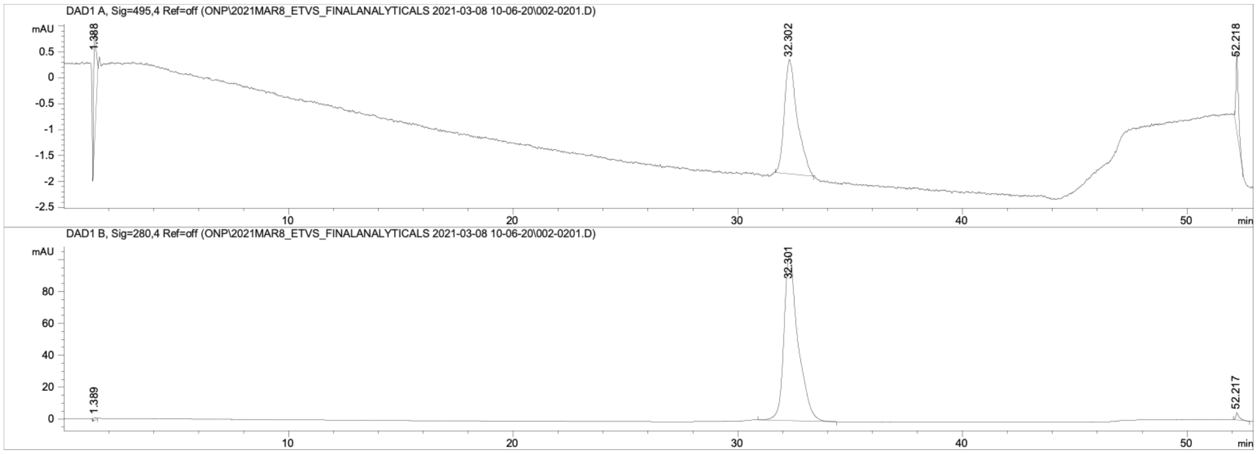


*Figure S32.* Analytical HPLC trace of **FITC-ETV1 (38-69)** monitored at 495 nm (top) and 280 nm (bottom).

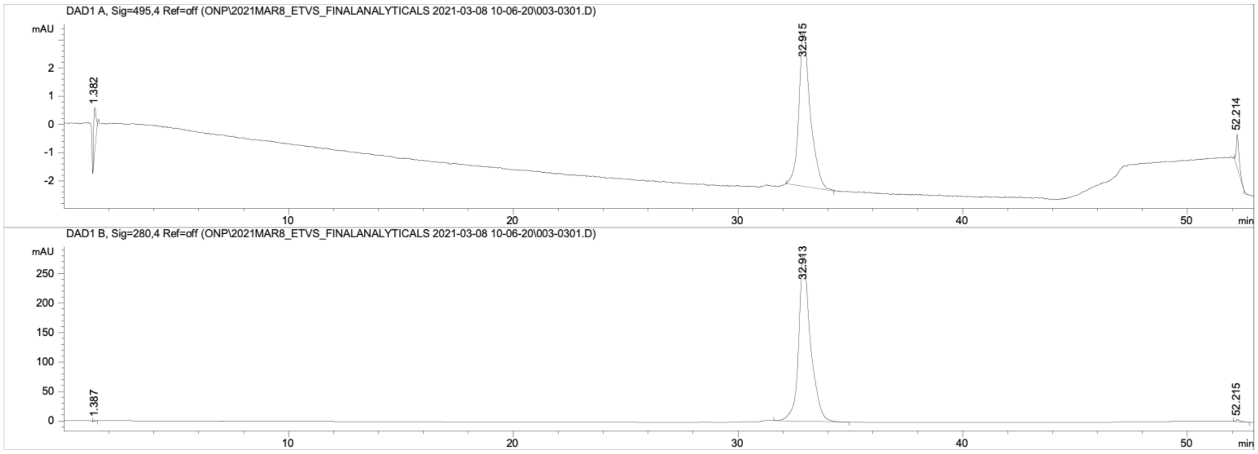


*Figure S33.* Analytical HPLC trace of **FITC-ETV4 (45-76)** monitored at 495 nm (top) and 280 nm (bottom).

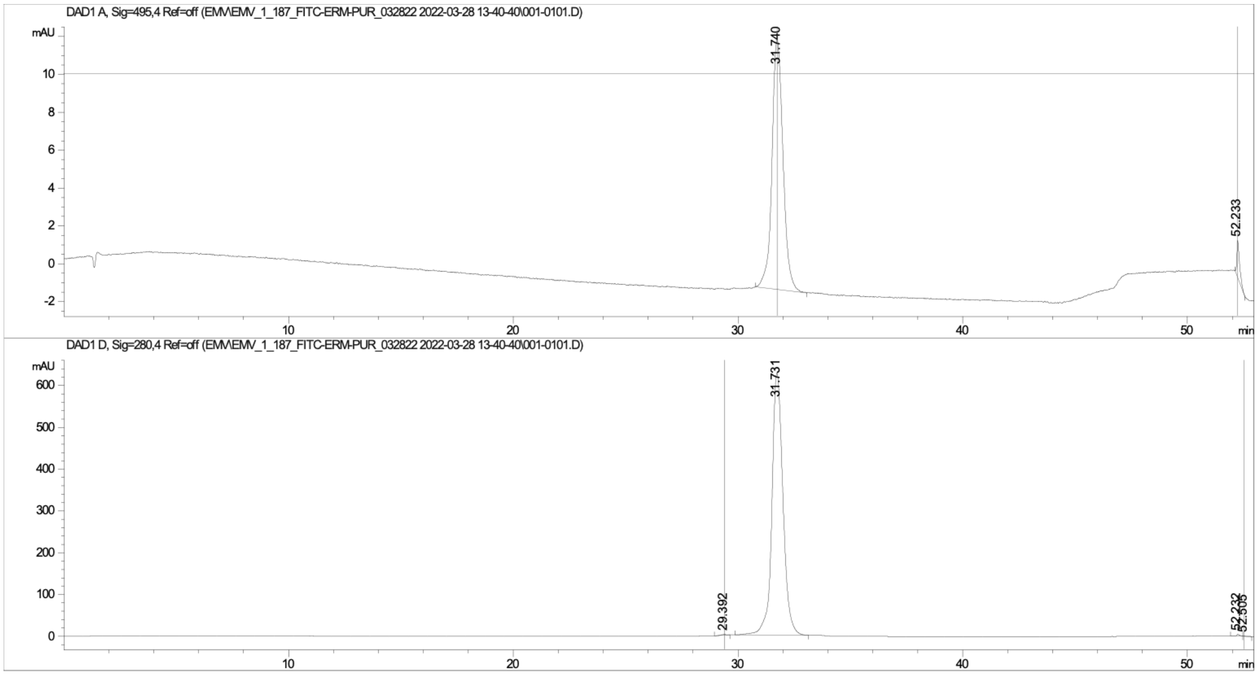


*Figure S34.* Analytical HPLC trace of **FITC-ETV5 (38-68)** monitored at 495 nm (top) and 280 nm (bottom).

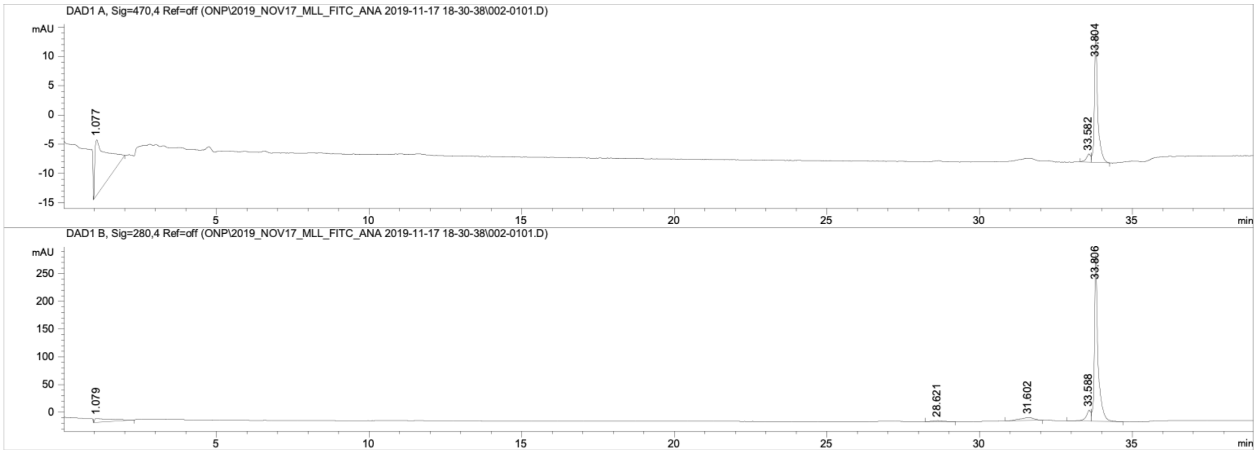


*Figure S35.* Analytical HPLC trace of **FITC-MLL (2840-2858)** monitored at 495 nm (top) and 280 nm (bottom).

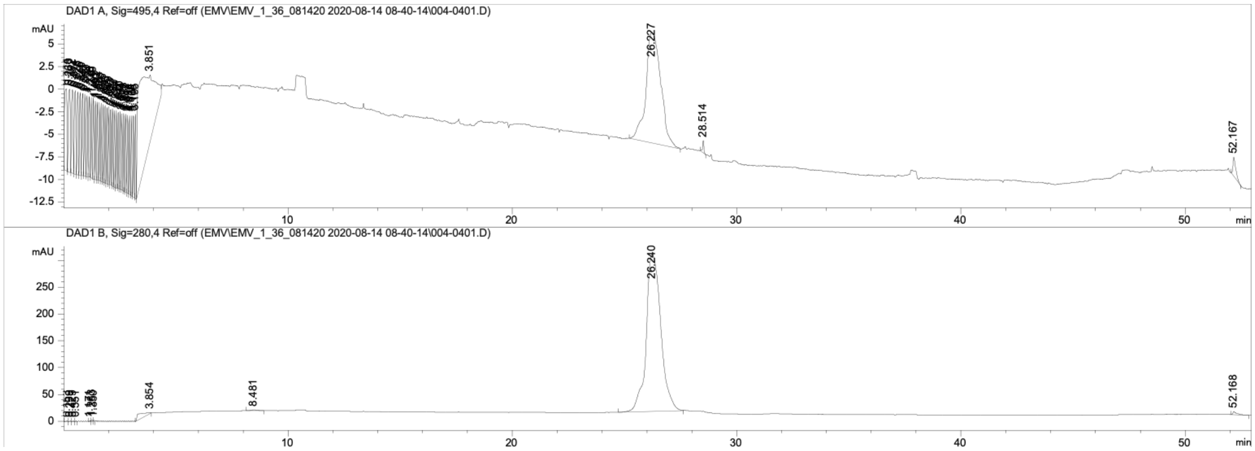


*Figure S36***.** Analytical HPLC trace of **FITC-Myb (219-316)** monitored at 495 nm (top) and 280 nm (bottom).

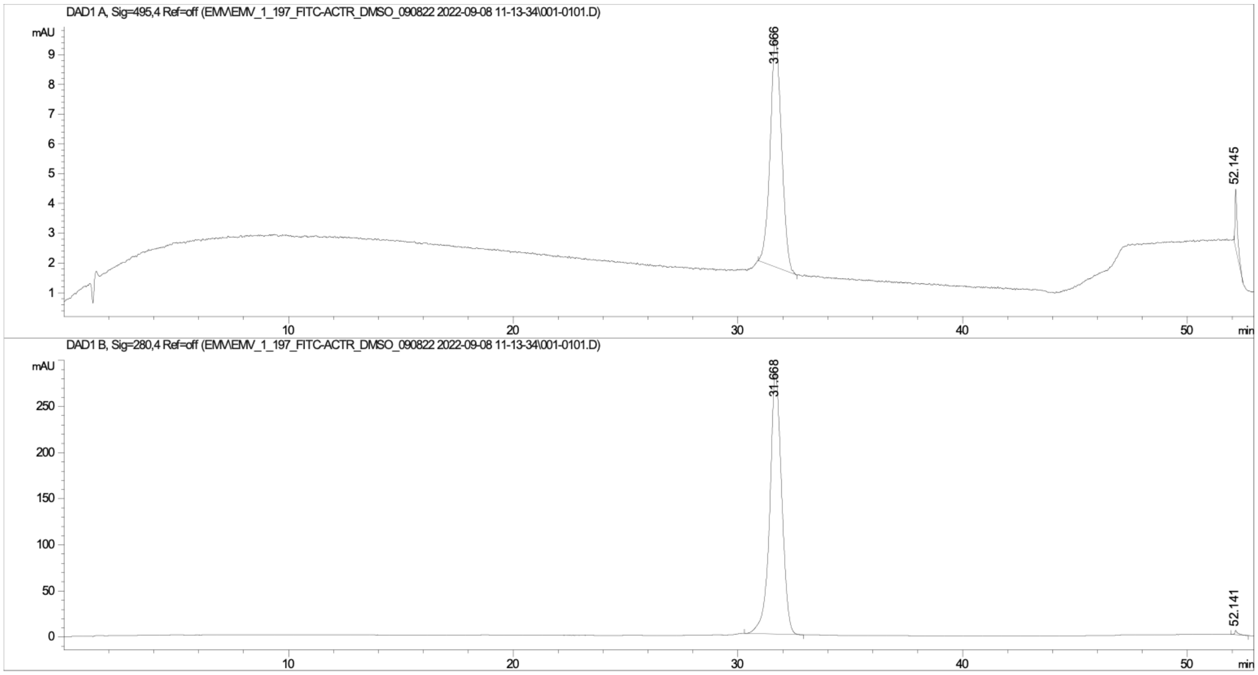


*Figure S37.* Analytical HPLC trace of **FITC-ACTR (1041-1088)** monitored at 495 nm (top) and 280 nm (bottom).

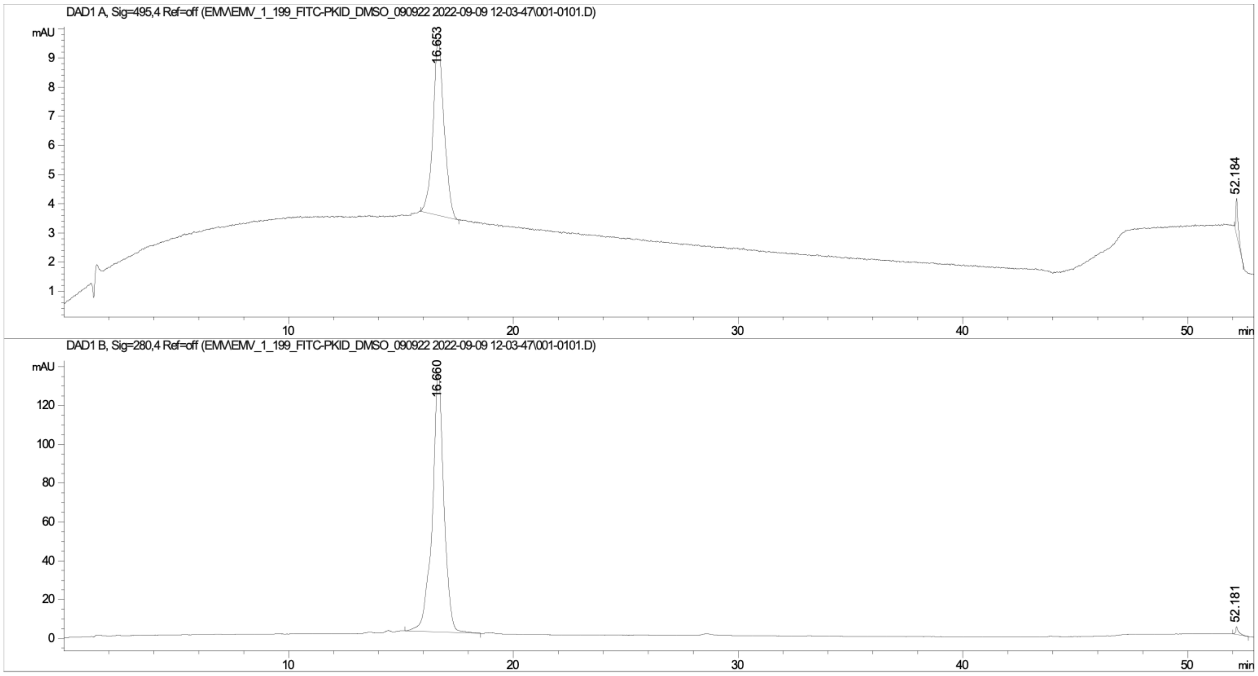


*Figure S38.* Analytical HPLC trace of **FITC-pKID (105-133)** monitored at 280 nm (top) and 495 nm (bottom).

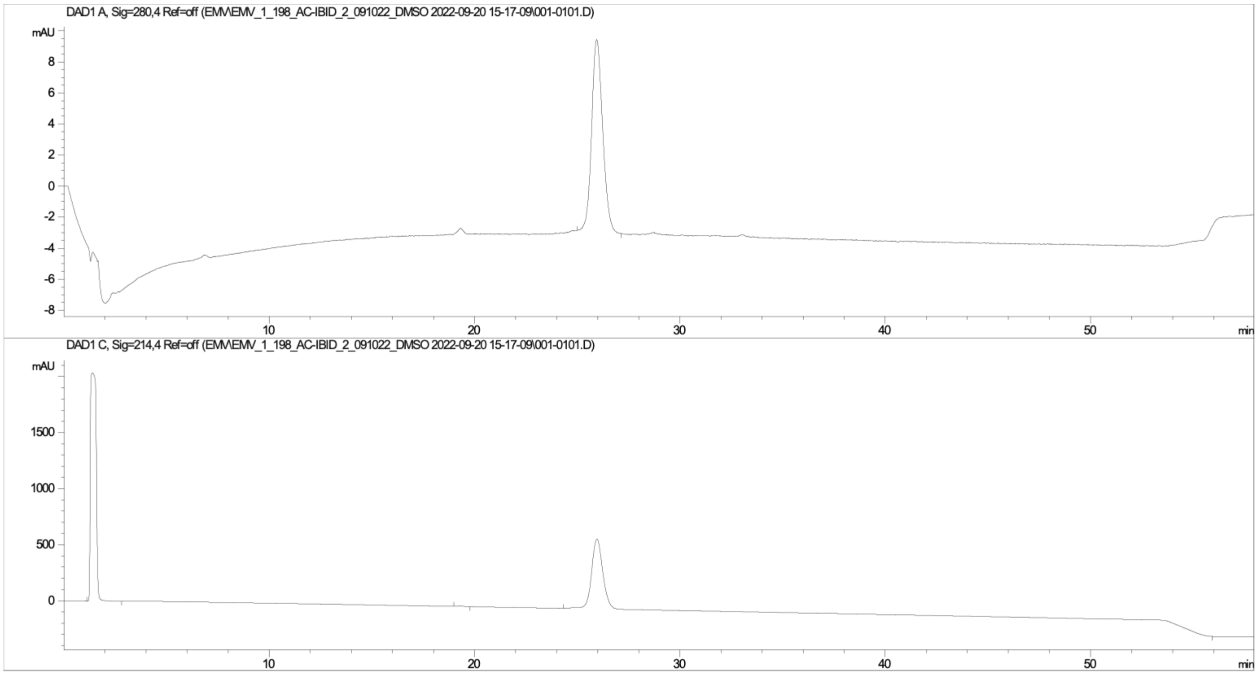


*Figure S39*. Analytical HPLC trace of Ac-IBiD (2063-2111) monitored at 280 nm (top) and 214 nm (bottom). Peak before 5 min. in 214 nm trace corresponds to solvent peak, DMSO, that was used to dissolve sample.

*High Resolution Mass Spectrometry of Lipopeptidomimetics*

34913-2 MS

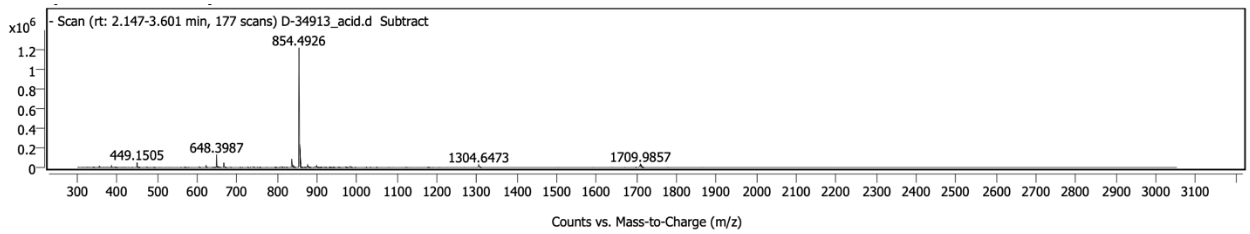


34913-2 Deconvolution

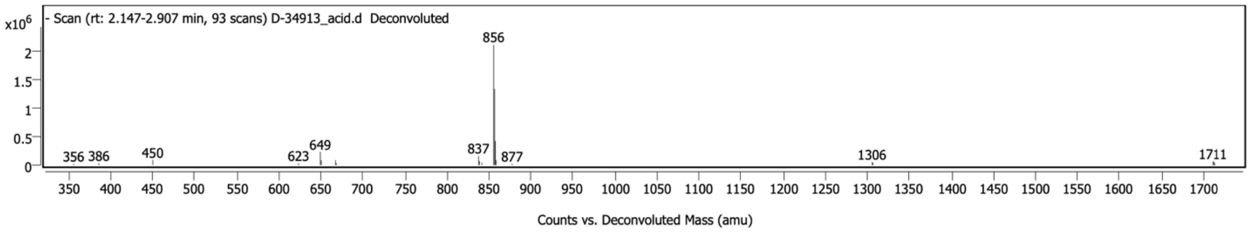


34913-3 MS

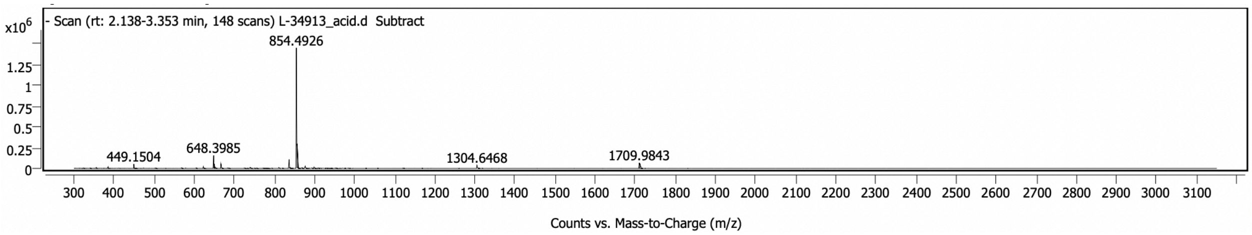


34913-3 Deconvolution

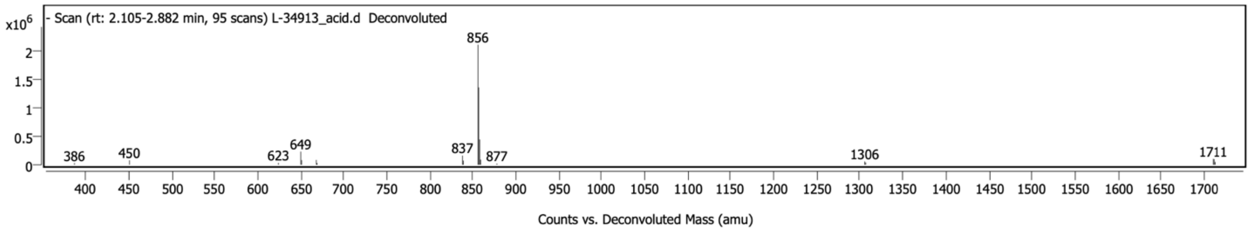


34913-4 MS

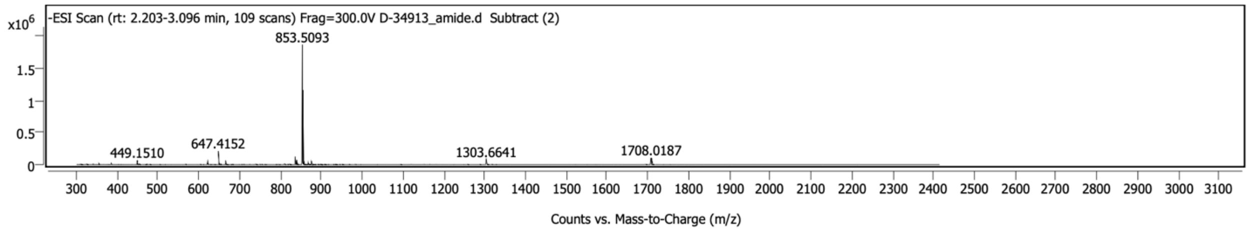

34913-4 Deconvolution

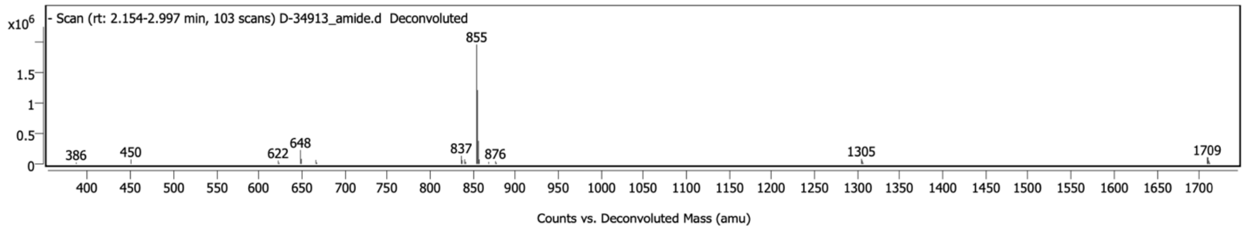


34913-5 MS

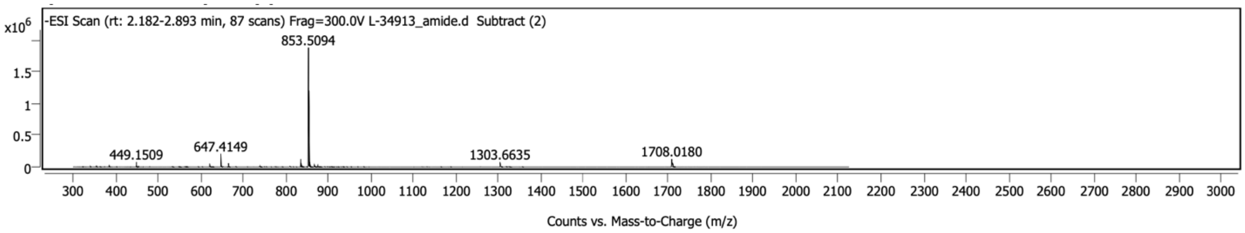


34913-5 Deconvolution

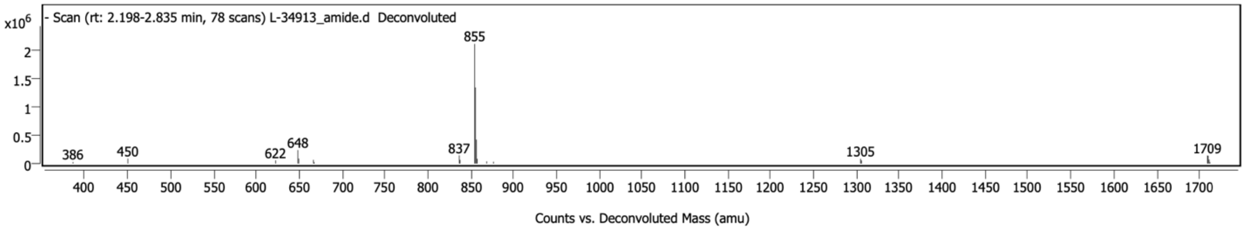


34913-6 MS

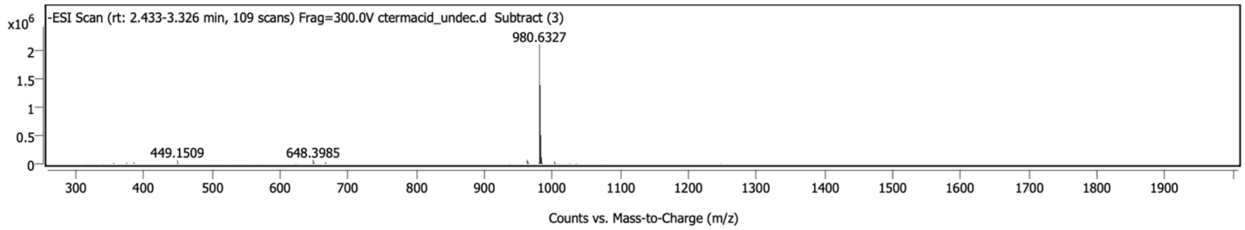


34913-6 Deconvolution

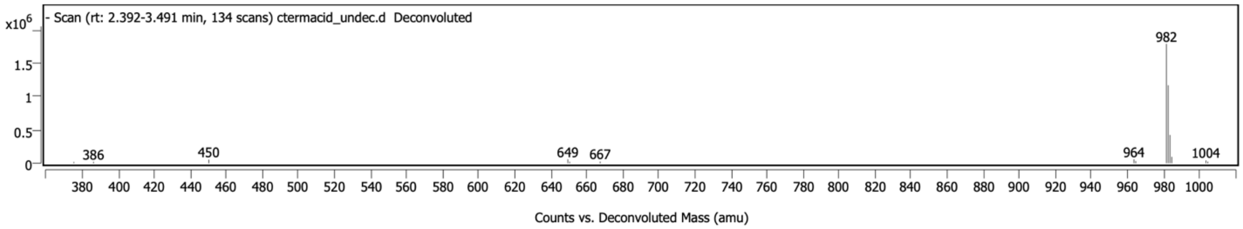

34913-7 MS

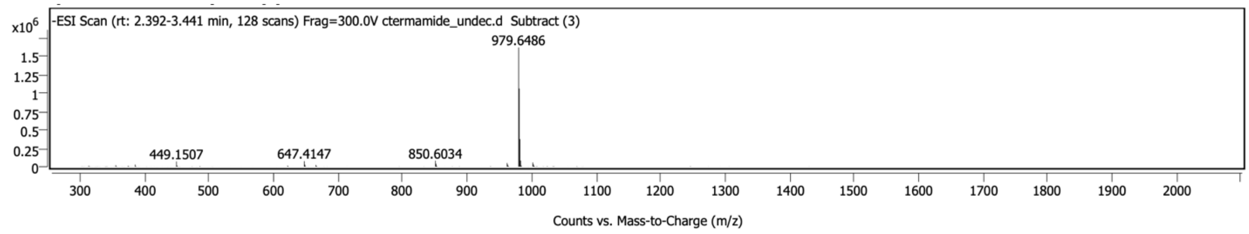


34913-7 Deconvolution

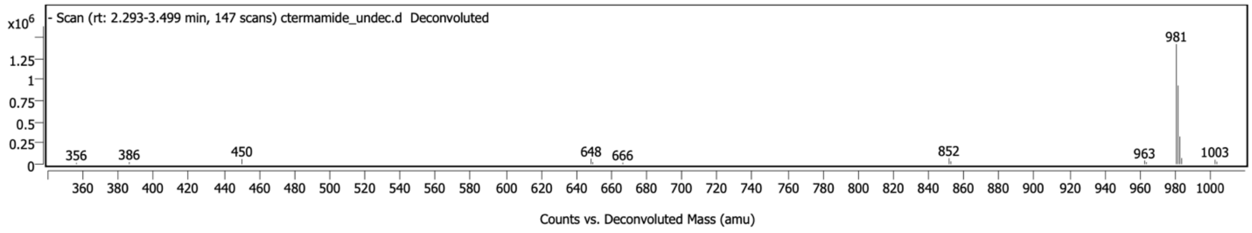


34913-8 MS

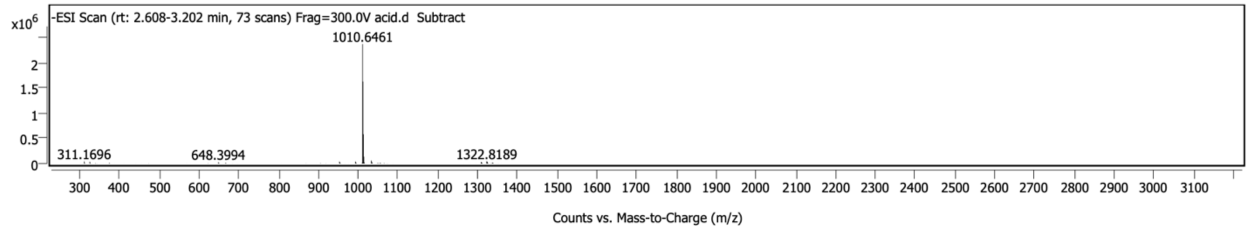


34913-8 Deconvolution

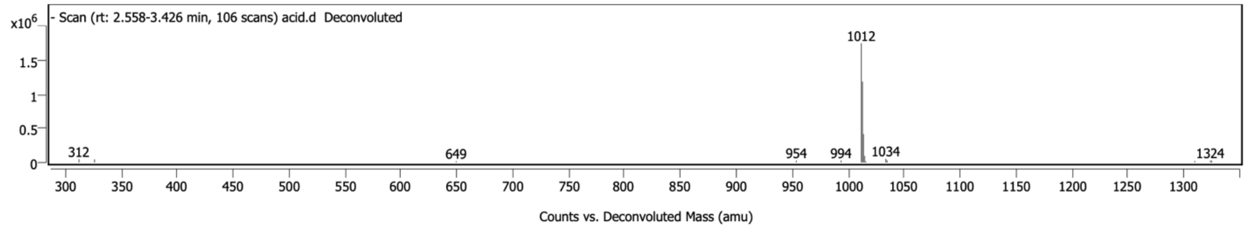


34913-9 MS

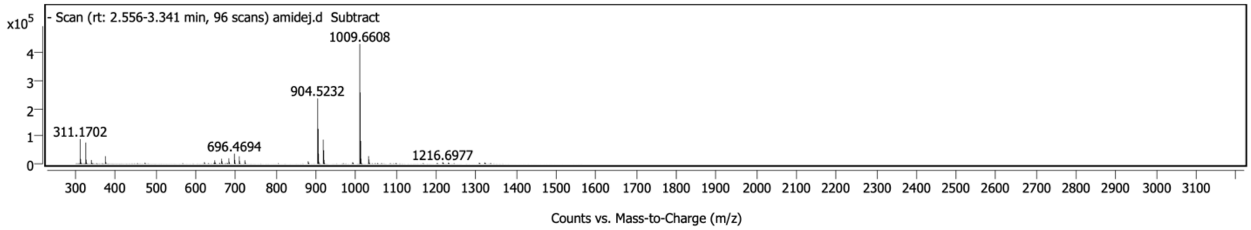

34913-9 Deconvolution

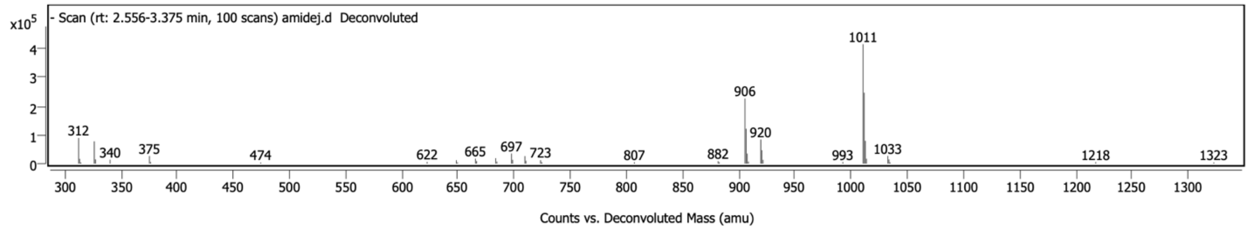


34913-10 MS

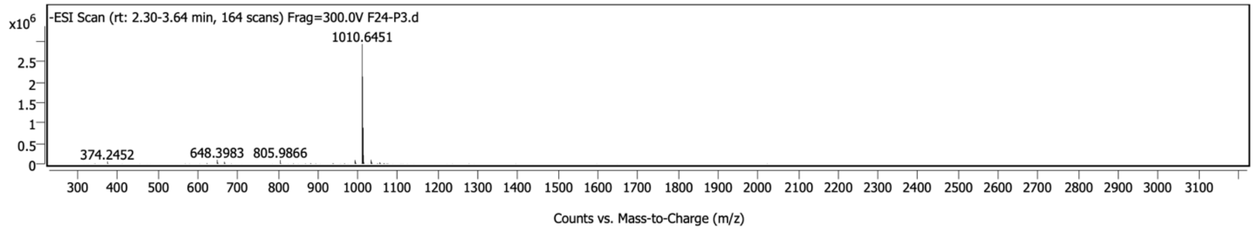


34913-10 Deconvolution

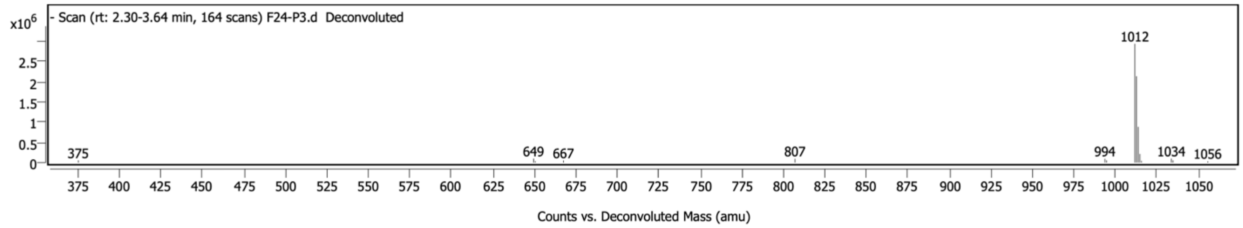


*Activator Peptide Mass Spectrometry*

FITC-ATF6α (38-75) MS

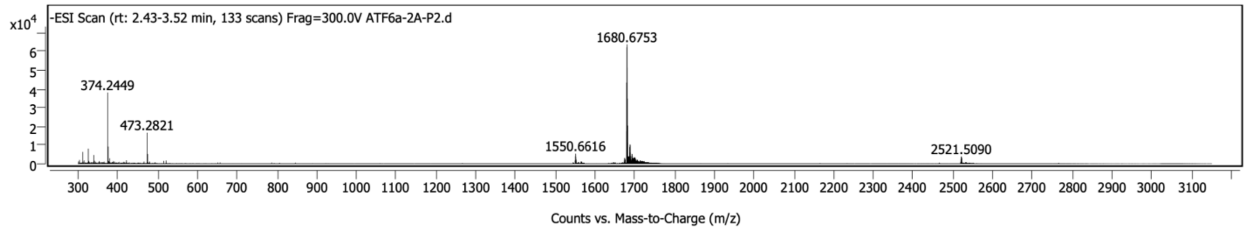


FITC-ATF6α (38-75) Deconvolution

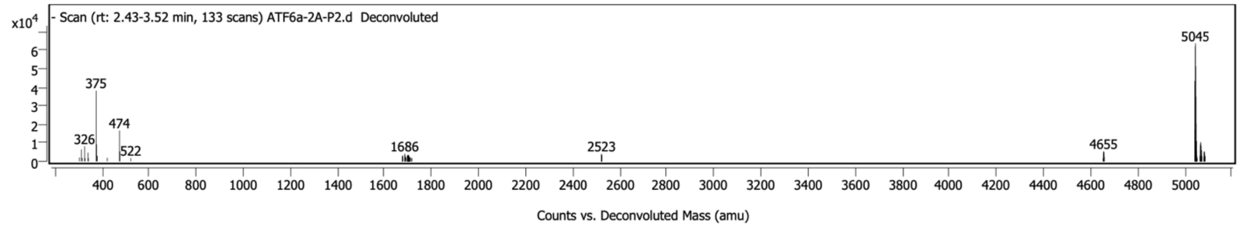

FITC-ETV1 (38-69) MS

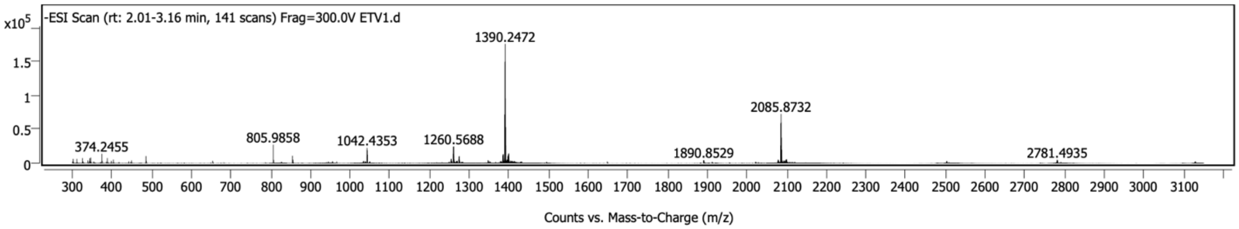


FITC-ETV1 (38-69) Deconvolution

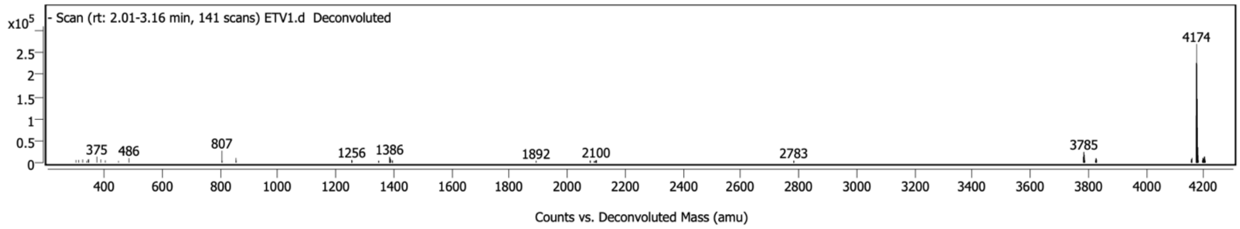


FITC-ETV4 (45-76) MS

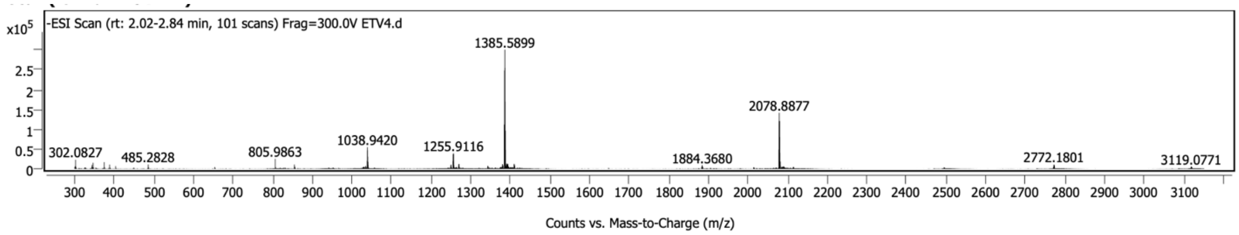


FITC-ETV4 (45-76) Deconvolution

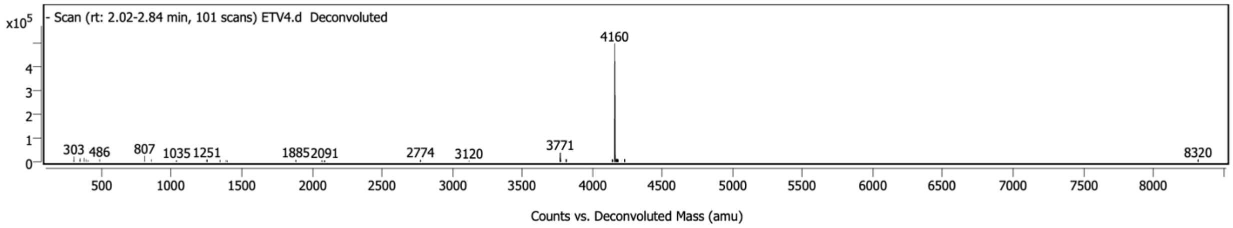


FITC-ETV5 (38-68) MS

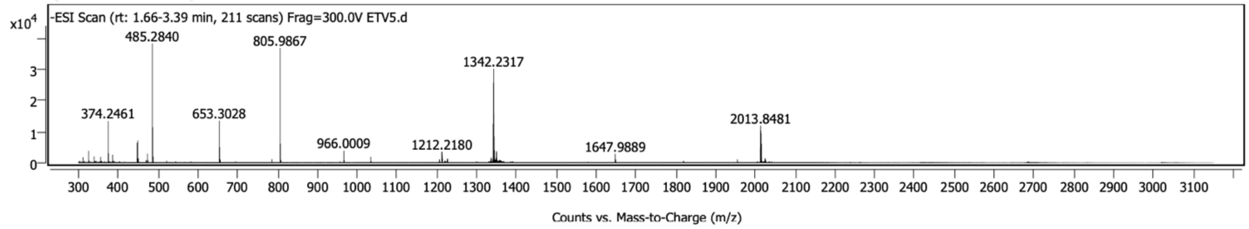

FITC-ETV5 (38-68) Deconvolution

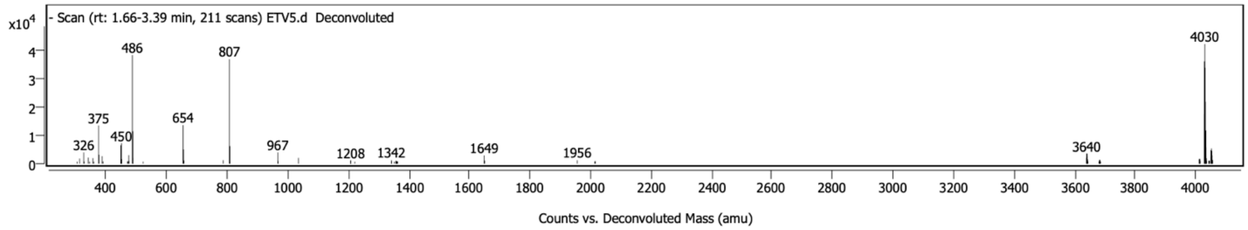


FITC-MLL (2840-2858) MS

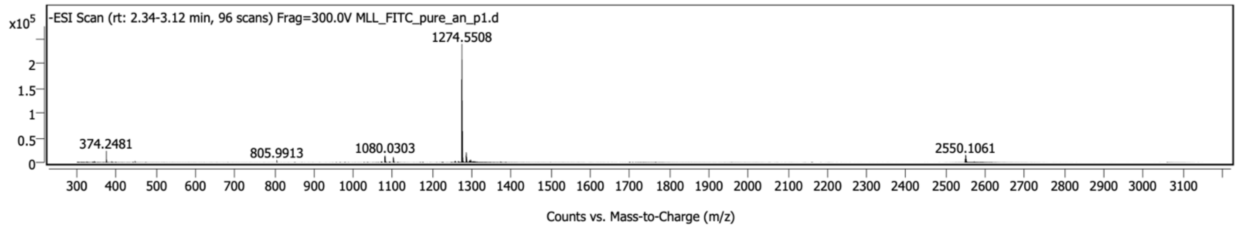


FITC-MLL (2840-2858) Deconvolution

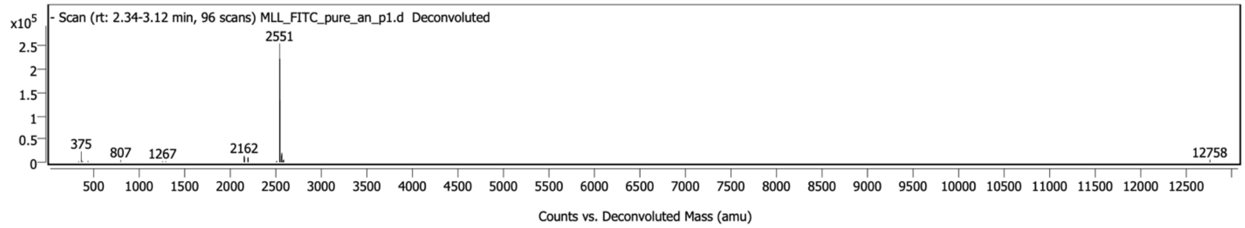


FITC-Myb (219-316) MS

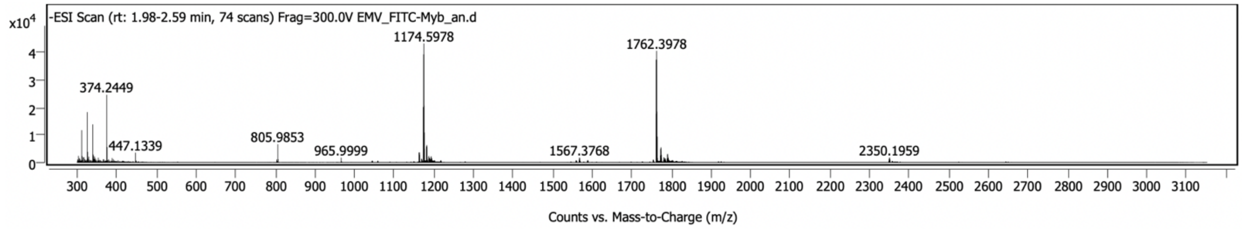


FITC-Myb (219-316) Deconvolution

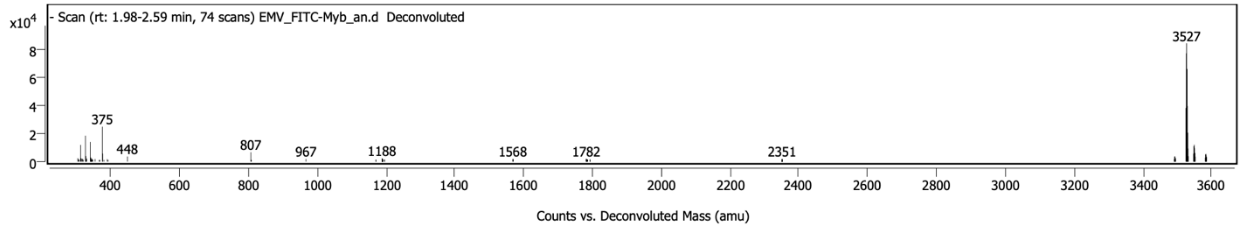


FITC-ACTR (1041-1088) MS

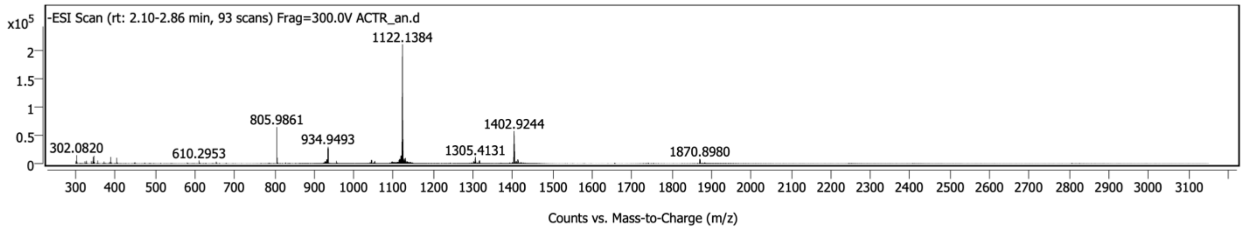

FITC-ACTR (1041-1088) Deconvolution

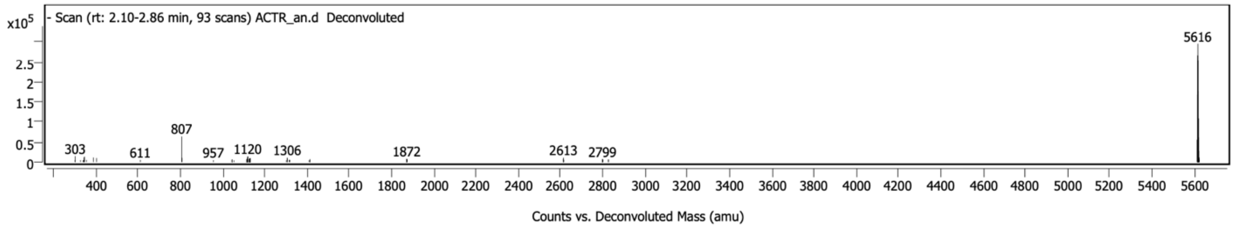


FITC-pKID (105-133) MS

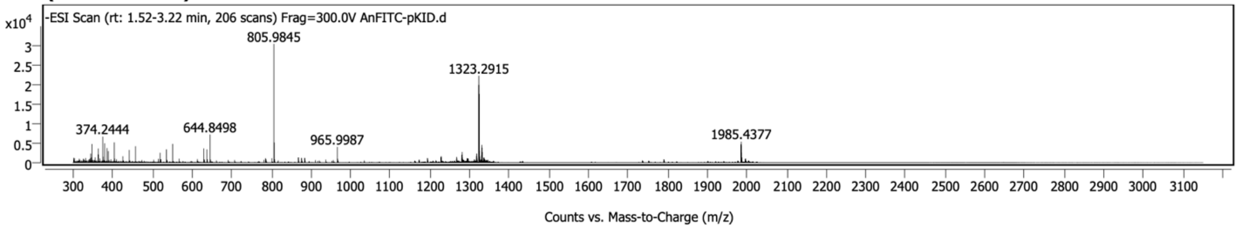


FITC-pKID (105-133) Deconvolution

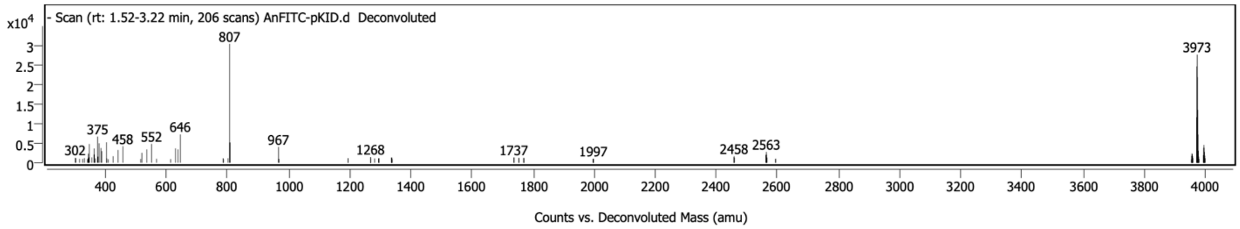


Ac-IBiD (2063-2111) MS

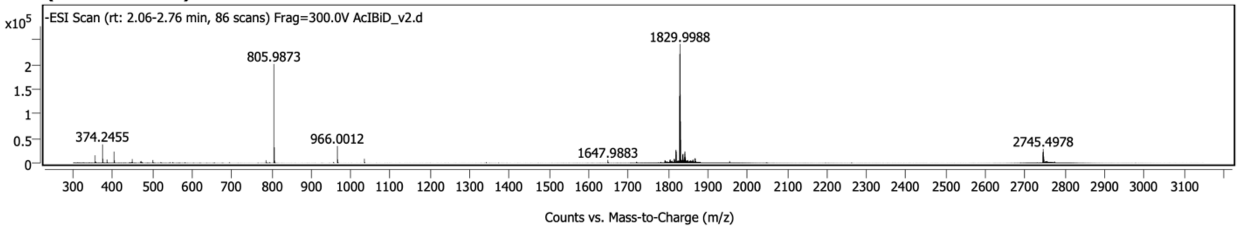


Ac-IBiD (2063-2111) Deconvolution

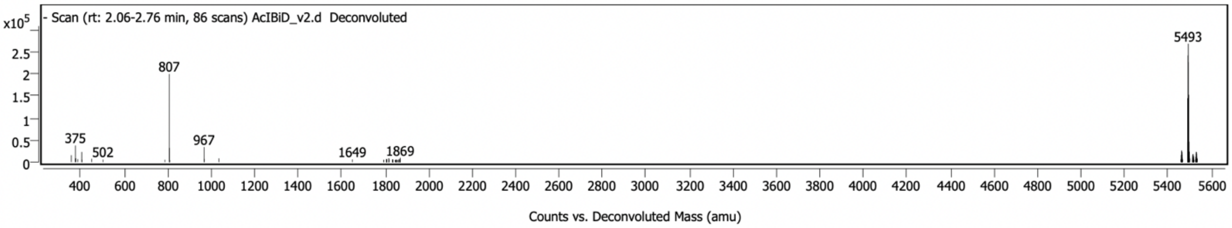

| Peptide | Structure | Exact Mass (M+1) | Observed Mass (M+1) |
|---|---|---|---|
| *34913-2* | 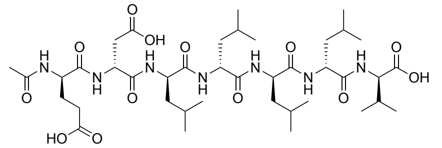 | 855.4953 | 855.5003 |
| *34913-3* | 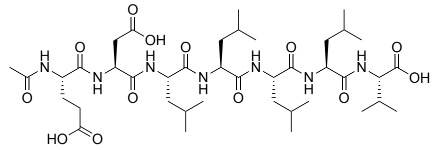 | 855.4953 | 855.5003 |
| *34913-4* | 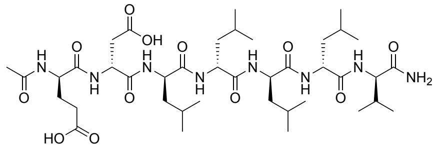 | 854.5113 | 854.5168 |
| *34913-5* | 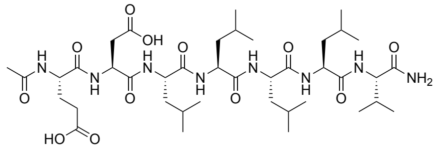 | 854.5113 | 854.5168 |
| *34913-6* | 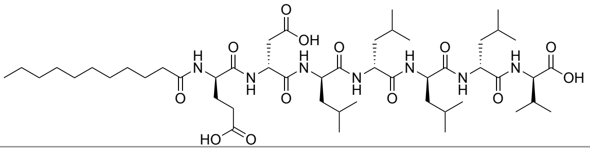 | 981.6362 | 981.6399 |
| *34913-7* | 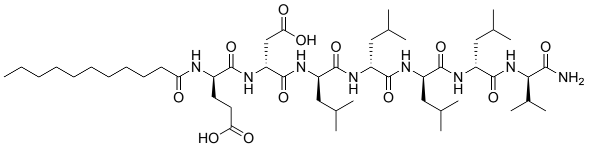 | 980.6522 | 980.6559 |
| *34913-8* | 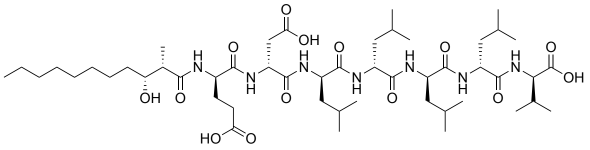 | 1011.6468 | 1011.6532 |
| *34913-9* | 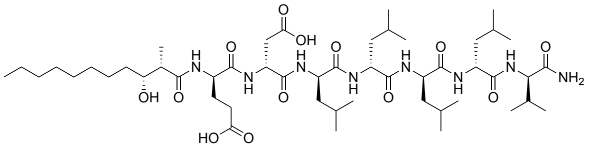 | 1010.6627 | 1010.6680 |
| *34913-10* | 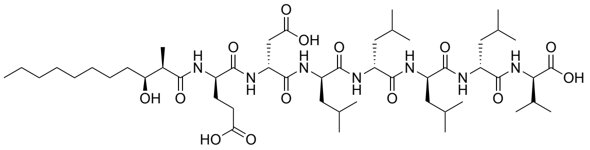 | 1011.6468 | 1011.6524 |

*Table S3.* Lipopeptide structures, expected masses based on structure and observed masses as determined on an Agilent LC/Q-TOF. Observed masses were obtained via deconvolution on an Agilent 6545 Q-TOF.

| Peptide | Exact Mass (M+1) | Observed Mass (M+1) |
|---|---|---|
| *FITC-ATF6α (38-75)* | 5045.0405 | 5045.0478 |
| *FITC-ETV1 (38-69)* | 4173.7598 | 4173.7637 |

| | | |
|---|---|---|
| *FITC-ETV4 (45-76)* | 4159.7846 | 4159.7915 |
| *FITC-ETV5 (38-68)* | 4029.7063 | 4029.7150 |
| *FITC-MLL (2840-2858)* | 2551.0935 | 2551.1160 |
| *FITC-Myb (219-316)* | 3526.81 | 3526.8126 |
| *FITC-ACTR (1041-1088)* | 5616.6928 | 5615.7288 |
| *FITC-pKID (105-133)* | 3972.8839 | 3972.8951 |
| *Ac-IBiD (2063-2111)* | 5493.9855 | 5493.0173 |

*Table S4.* FITC-labelled and acetylated transcriptional activator domain peptides, expected masses based on structure, and observed masses as determined on an Agilent LC/QTOF. Observed masses were obtained via deconvolution on an Agilent 6546 Q-TOF.
